# Polyethylene terephthalate (PET) primary degradation products affect c-di-GMP-, cAMP-signaling and quorum sensing (QS) in *Vibrio gazogenes* DSM 21264

**DOI:** 10.1101/2025.01.28.635239

**Authors:** Lena Preuss, Malik Alawi, Albert Dumnitch, Ly Trinh, Wolfgang Maison, Nils Burmeister, Anja Poehlein, Rolf Daniel, Christel Vollstedt, Wolfgang R. Streit

**Affiliations:** Department of Microbiology and Biotechnology, University of Hamburg, Ohnhorststr.18, D-22609 Hamburg. Germany; Bioinformatics Core, UKE Hamburg, Martinitstr. 52. D-20246 Hamburg, Germany; Department of Chemistry, University of Hamburg, Bundesstr. 45, D-20146 Hamburg. Germany; Department of Genomic and Applied Microbiology, Georg-August University of Göttingen, Grisebachstr. 8, D-37077 Göttingen, Germany

**Keywords:** *Vibrio* sp. biofilms on plastics, plastisphere, polyethylene terephthalate (PET), bis(2-hydroxyethyl) terephthalate, mono-(2-hydroxyethyl) terephthalate (MHET), phages

## Abstract

Global plastic pollution in oceans and estuaries is increasing rapidly and it’s well known that bacteria colonize plastic particles of all sizes. *Vibri*o spp. are frequently found as part of the plastisphere. We recently showed that *Vibrio gazogenes* DSM 21264 harbors a promiscuous esterase designated PET6. We now provide evidence that the *pet6* gene is expressed under a wide range of environmental conditions in its native host. However, in PET- and PE-grown biofilms the *pet6* gene expression was not affected by the type of surface. The *pet6* transcription was sufficient to allow enzyme production and release of µM amounts of mono-(2-hydroxyethyl) terephthalate (MHET) and terephthalic acid (TPA) already after 24 hours of incubation on PET foil. Notably, the highest *pet6* gene transcription was observed in planktonic lifestyle in the presence of bis(2-hydroxyethyl) terephthalate (BHET) one of the primary degradation products of PET. BHET was further hydrolyzed by PET6 and UlaG, a lactonase that had not been known to be involved in BHET degradation. Elevated concentrations of BHET affected the major signaling circuits involved in bacterial quorum sensing (QS), c-di-GMP and cAMP-CRP signaling. This resulted in failure to form biofilms, synthesis of the red pigment prodigiosin and altered colony morphologies. While BHET had a very wide impact, TPA interfered mainly with the bacterial QS by attenuating the expression of the CAI-I autoinducer synthase gene. These observations imply a potential role of BHET and TPA as nutritional signals in *Vibrio gazogenes* and that may affect its growth and survival in the plastisphere.

**IMPORTANCE:** This study provides first evidence that *Vibrio gazogenes* DSM 21264 secretes an active PET hydrolase and degrades the polymer using PET6 when growing in biofilms on foils and microplastic particles. The study further provides evidence that the primary PET degradation products BHET and TPA may have a profound impact on the global QS, c-di-GMP and cAMP-CRP signaling of *V. gazogenes* and its capability to colonize plastic particles in the marine environment.

## INTRODUCTION

Global plastic pollution has reached an alarming and unprecedented level in the recent decade. Recently, it was estimated that a minimum of 8-10 million tons of plastic is entering the oceans annually [1], [2], [3], [4], [5]. While most of the plastic enters the environment as larger floating fragments, weathering results in the production of smaller particles (micro-, nano- and pico-plastics) with sizes less than 5 mm in diameter [6], [7]. These particles and the additives contained in them are believed to have negative impact on all ecological niches and their biodiversity [8], [9]. Ultimately, they will also affect human health and wellbeing [10].

The majority of the fossil-fuel based and synthetic polymers which end up in the environment are polyethylene (PE), polypropylene (PP), polyvinylchloride (PVC), polyethylene terephthalate (PET), polyurethane (PUR), polystyrene (PS) and polyamide (PA) only being produced at lower quantities [11]. While for most of these polymers no enzymes and microorganisms are known to degrade them the degradation of PET is well understood (www.PAZy.eu) [12]. PET is degraded by promiscuous and secreted esterases, lipases or cutinases that all have a wide substrate specificity and are members of the E.C classes 3.1.1.-[11], [13], [14]. In addition to these studies recently much effort has been made to analyze and characterize the phylogenetic makeup of microbial communities associated with plastic particles found in the marine environment. The microorganisms colonizing plastic particles (i.e. the plastic microbiota) have been termed ‘plastisphere’ [15], [16], [17], [18]. These studies indicate that growth on and the initial colonization of plastic surfaces depends on the chemical and the physical properties of the various polymers. Thereby, the surface charge, roughness and the different chemical compositions of additives play key roles during the attachment and growth of the plastic colonizing microbiota [19], [16].

Notably, the simple colonization does not imply any biodegradation [6]. The microbial communities observed are highly diverse in their phylogeny and bacteria affiliated with the genus *Vibrio* are often observed on plastic particles [20] [18] [21] [22] [23] [24]. *Vibrio* species are ubiquitously occurring gram-negative bacteria that are mainly found in marine environments where they are considered to be key players in carbon and nitrogen cycling [25] [26]. There are more than 100 known *Vibrio* species of which few are either human or fish pathogens (e.g. *Vibrio cholerae*, *Vibrio parahaemolyticus, Vibrio alginolyticus, Vibrio vulnificus* and others) [27] [28] [29].

Recently, we have shown that the marine organism *V. gazogenes* DSM 21264 (from here on DSM 21264) harbors a gene encoding a promiscuous esterase designated PET6 in its 4.6 Mbp genome [30] [31]. DSM 21264 (synonym PB1, AATCC 29988) is a gram-negative and non-pathogenic bacterium. It was isolated from sulfide-containing mud collected from a saltwater marsh and produces a red pigment, prodigiosin [32]. The species is globally occurring and typically found in estuaries. The *pet6* gene is encoded by ORF AAC977_05355. The heterologous expressed protein hydrolyzes amorphous PET and other substrates including short chain fatty acid esters [30] [31] and its activity is salt-dependent. PET hydrolysis catalyzed by recombinant PET6 is, however, rather low compared to other known enzymes used in industrial processes like the well-characterized IsPETase or LCC [31]. While much information was gathered on the enzyme structure and biochemical function of PET6, it was not known under which conditions the *pet6* gene is transcribed in its native host and if the native enzyme would result in PET degradation in the environment.

Within this manuscript we provide first evidence that the *pet6* gene is transcribed at low levels under various environmental conditions. The *pet6* gene expression is, however, not affected by the presence of PET, PE foil or PET powder.

Instead, BHET, a primary PET degradation product, affects *pet6* transcription at mM concentrations. Further, our data imply that BHET is a nutritional signal affecting the c-di-GMP, the cAMP-CRP and QS-dependent signaling pathways in DSM 21264. These three signaling pathways are essential to lifestyle transitions from motility, attaching to a surface and forming biofilms in the plastisphere.

## RESULTS

### DSM 21264 forms patchy biofilms on PE and PET independent from the surface and releases µM amounts of MHET and TPA

Since we had earlier shown that DSM 21264 produces a PET-active hydrolase, designated PET6 [31] [30], we wanted to know if the organism forms biofilms on PET foil and actively degrades PET. Further, we asked if the *pet6* gene is expressed at significant levels. Therefore, we inoculated DSM 21264 in artificial seawater medium (ASWM) with PET or PE foil for biofilm formation and potential degradation. Since no PE-degradative genes have been reported in gram-negative bacteria PE-foil was used as a control. In these biofilm experiments cells attached in general at very low frequencies and only thin single cell layer biofilms were formed after 10 and 180 days (FIGURE 1 & S1). Therefore, we used ASWM supplemented with tryptone and yeast extract 1 % (w/v) in further tests to promote growth and biofilm formation. Under these conditions DSM 21264 attached within few hours and formed patchy biofilms on both plastic surfaces, whereby the biofilms formed on PE were less patchy than those on PET (FIGURE 1). Biofilms formed on PE and PET had an average thickness of 2-3 μm equaling one cell layer (FIGURE 1).

**FIGURE 1:**
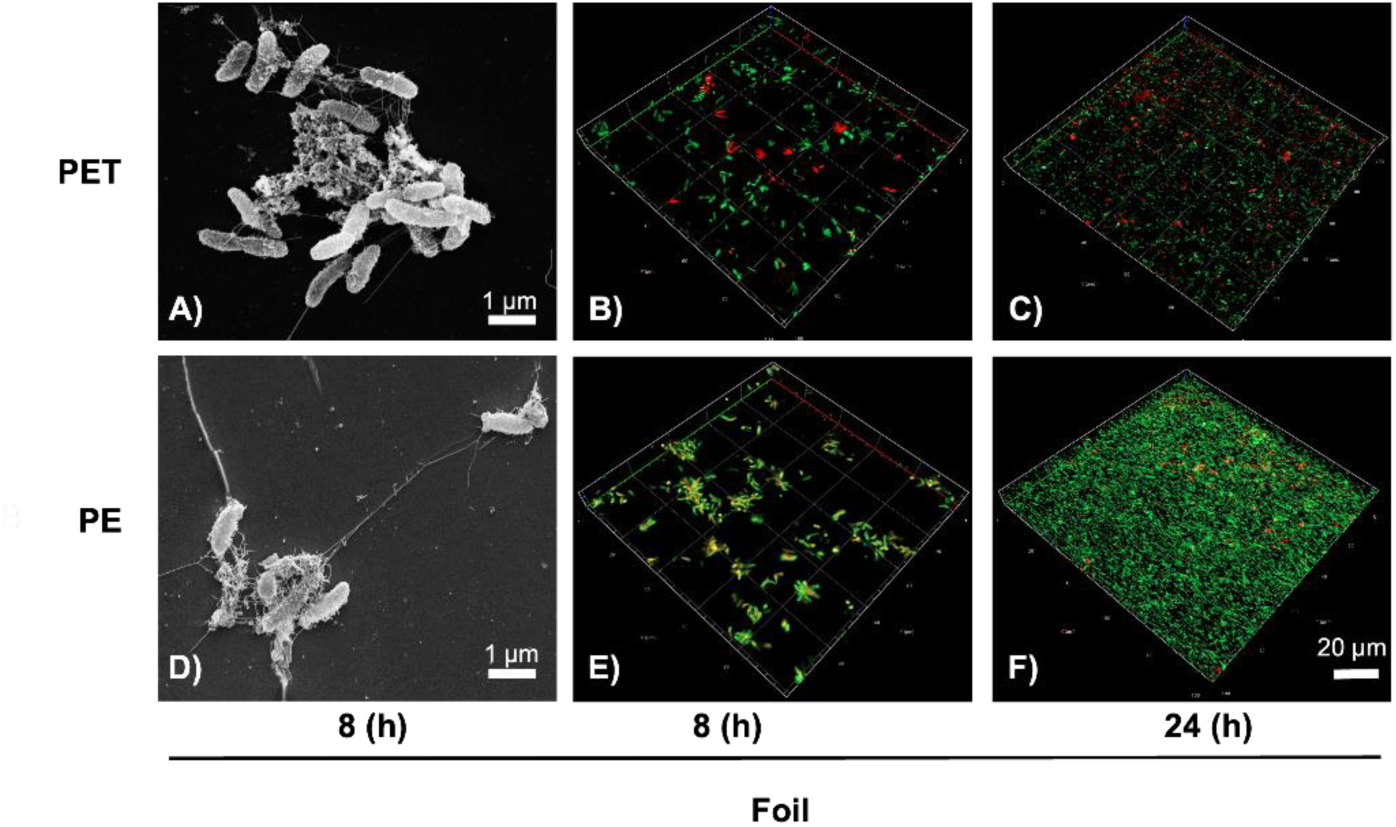
Scanning electron microscope (SEM) **(A, D)** and confocal laser scanning microscope (CLSM) images **(B, C, E, F)** of *V. gazogenes* DSM 21246 biofilms grown on PET and PE foil. SEM images and CLSM images **(C & D)** show first attachment of cells after 8 hours of incubation in ASW medium at 22 °C of incubation on PET foil **(A & C)** and PE foil **(B & D)**. CLSM images **(E & F)** show biofilm formation of DSM 21264 on PET foil **(E)** and PE foil **(F)** after 24 hours of incubation. Cells were stained using LIVE/DEAD stain.

SEM images showed that the cells on PET and PE were interconnected by fiber- and net-like structures with a length of 4-10 μm. However, the net-like structures were more prominent on PE surfaces (FIGURE 1). Additional tests with air plasma-treated PET and PE foil were conducted [33]. Air plasma treatment leads to the formation of oxygen-functionalities and thus an increase in surface polarity. It typically gives a potential degradative enzyme better access to the polymer fibers [34]. However, in these tests only minor differences were observed with respect to biofilm formation (FIGURES 1 & S1). Notably, when we grew DSM 21264 on PET foil or powder, we detected µM amounts of MHET and TPA in the biofilm and culture supernatants already after 24 hours (FIGURE 2).

**FIGURE 2:**
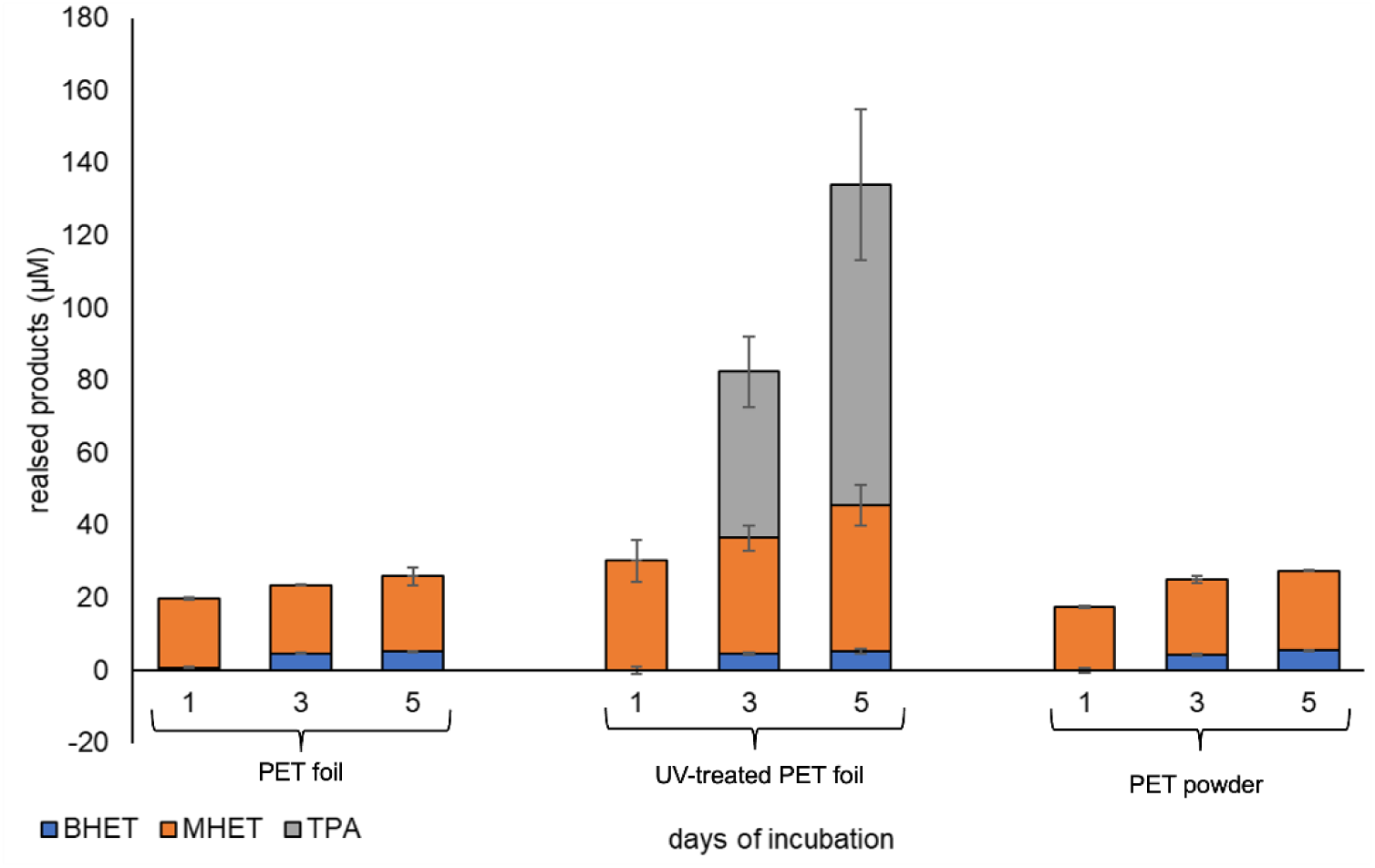
PET degradation products observed in supernatants of DSM 21264 biofilms grown on PET foil (non-treated), UV-treated foil and PET powder. DSM 21264 was grown in ASWM medium with reduced carbon source. Supernatants were collected after 1, 3 and 5 days and the degradation products were measured using UHPLC and as described in the Material and Methods section. Data were normalized and corrected to *E. coli* DH5α growing on PET foil and powder. Data are mean values of three independent replications and bars represent the standard deviations.

### Transcriptome analysis identifies keys to PET6 gene expression

Based on the above-made observations, we further asked if the *pet6* gene (AAC977_05355) was expressed and if surface attachment to PET or PE resulted in major transcriptional changes. To address these questions, we mainly used transcriptome studies with the goal to obtain an insight into the overall gene expression pattern of DSM 21264 in response to PET or other substrates during life in biofilms and planktonic cultures. In total we analyzed 14 different environmental parameters and/or carbon sources. Each experiment was repeated 3 times resulting in 42 RNAseq data sets which have been submitted to the European Nucleotide Archive (ENA). They are publicly available under accession PRJEB80907. Obtained reads were mapped against the genome of DSM 21264. For this purpose, we established a high-quality genome sequence of DSM 21264 that encompasses two chromosomes coding for 4,159 genes. Chromosome 1 (CP151640) codes for 3,121 genes and chromosome 2 (CP151641) for 1,038 genes. The newly established genome sequence is available under accession numbers CP151640 and CP151641 at NCBI/GenBank. An overview of the data sets reads obtained, the percentage of reads mapped to the genome and other parameters together with the level of *pet6* gene expression is given in TABLE S2 and FIGURE 3.

**FIGURE 3:**
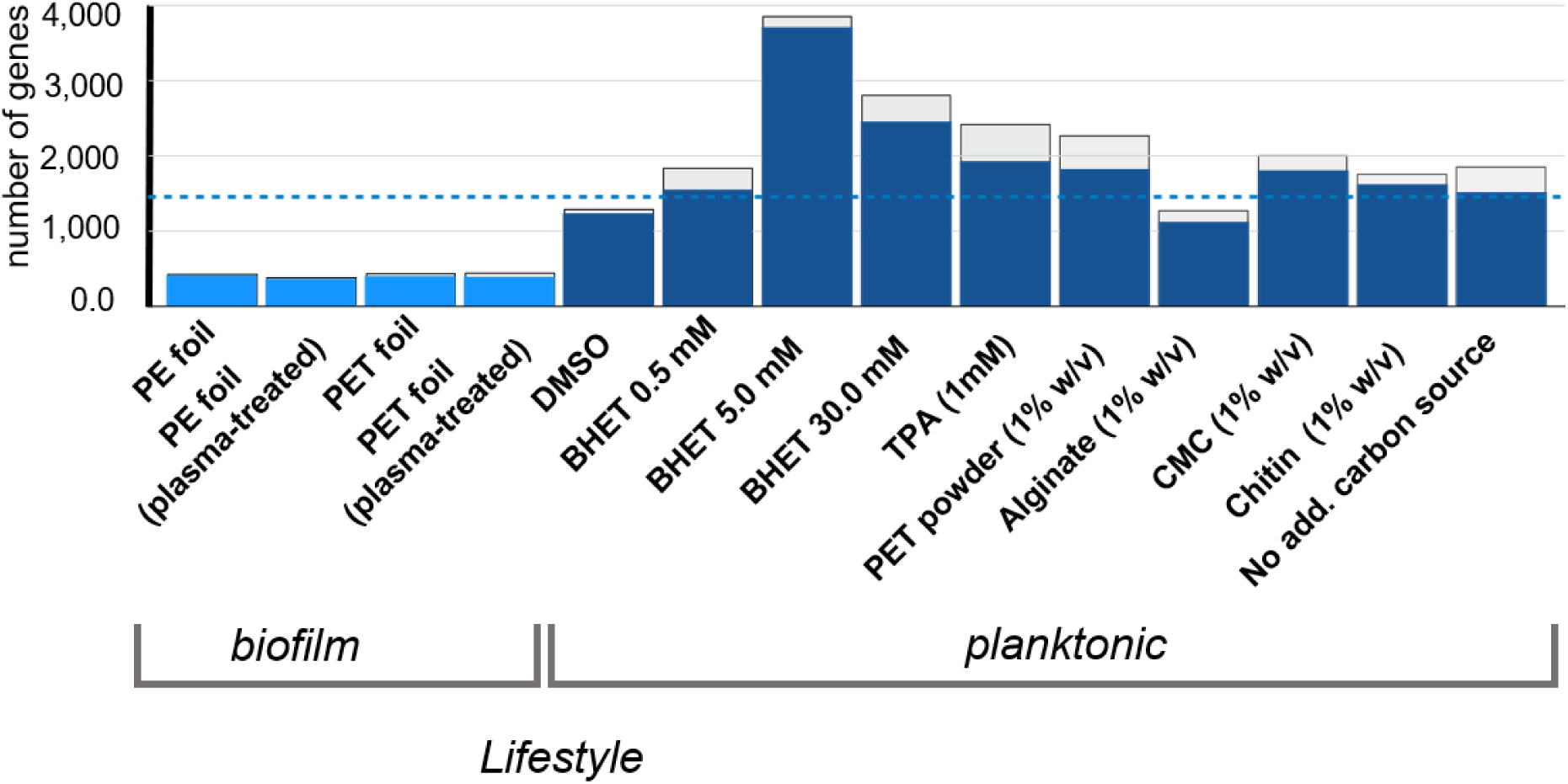
Transcription level of the *pet6* gene in relation to all 4,117 genes transcribed in DSM 21264. Blue bars indicate ranking of *pet6* depending on all genes transcribed. Standard deviations are indicated as light grey bars on top of the colored bars. The blue dotted line indicates the mean transcription level of all 4,117 genes in all 42 transcriptomes. Each data set represents the mean data from three independent biological replicates. Ribosomal, rRNA and tRNA genes were not included in the analyses.

### PET6 is weakly expressed in biofilms grown on PET and PE foils

When we compared PET-grown biofilms versus PE-grown biofilms, the expression *of pet6* was low and ranked around 25 % of the weakest transcribed genes. FIGURE 3 summarizes the level of expression of *pet6* under the biofilm conditions tested and in relation to the genome-wide expression of all genes. Plasma-treatment of the PET or PE foil had no significant impact on the low level of *pet6* gene expression (TABLE S1 and FIGURE 3). This observation is in line with the slow degradation of the polymer in biofilm cultures and as described above (FIGURE 2). In addition, we noticed, that in all biofilm experiments the transcription of *ompU* was among the highest transcribed genes implying that OmpU possibly is involved in surface colonization. Furthermore, only four genes were differentially expressed in all biofilm studies (TABLE S1 and S3). Highest log2-foldchanges were detected for a glutathione peroxidase and an alkyl hydroxide peroxidase, both involved in hydrogen peroxide scavenging [35]. In addition, the *adhE* gene, which is involved in ethylene glycol (EG) metabolism was significantly upregulated in PET biofilms but not in PE-grown biofilms (TABLE S1). To further verify the increased expression of *adhE,* a reporter fusion was constructed fusing the promoter with the amcyan gene in pBBR1-MCS-1 and mobilized into DSM 21264 (TABLE 1). Using the *P_adhE::amcyan* transcriptional reporter gene fusion in DSM 21264 we were able to verify that *adhE* was 2-fold upregulated in the presence of 5 mM EG (FIGURE S5).

**TABLE 1:**
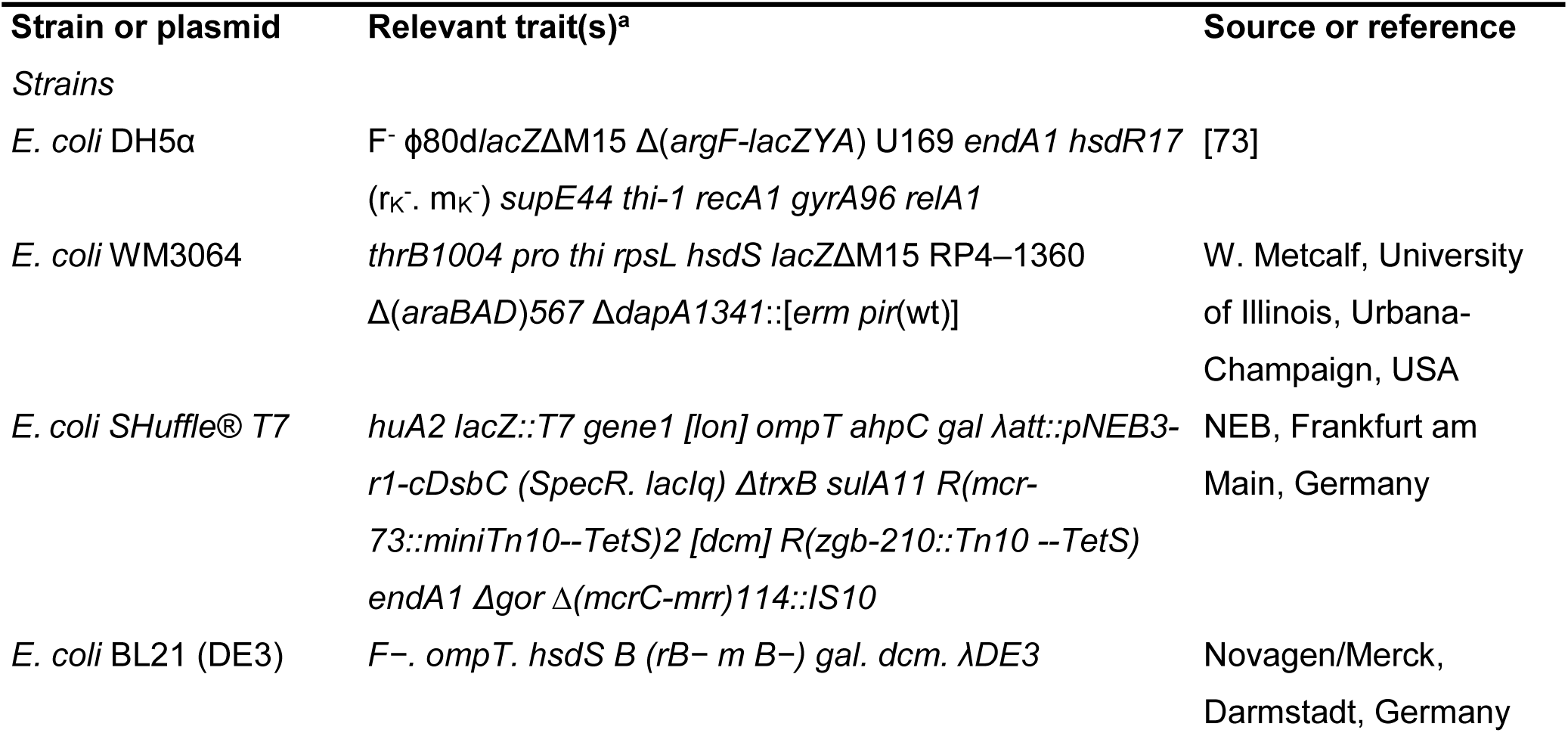
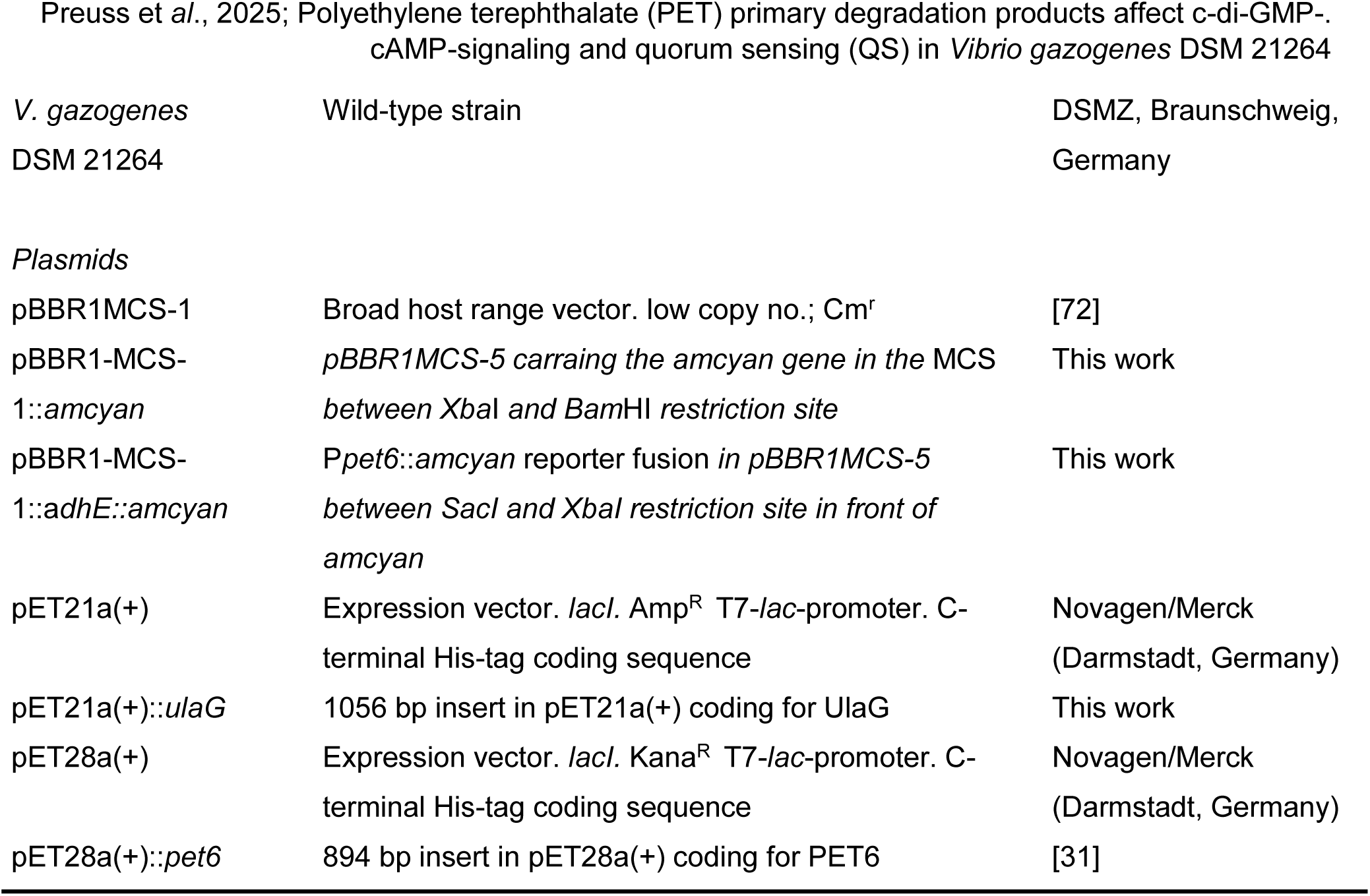
Bacterial strains and plasmids used in this study.

Since PET and PE are no natural substrate for bacteria, we further asked if other natural polymers would stimulate *pet6* gene expression. Therefore, we added alginate, chitin and carboxymethyl cellulose (CMC) (1 % w/v) to planktonic cultures and assayed the transcriptomes after 24 hours of aerobic growth. As expected in the planktonic cultures a largely different pattern of gene transcription was observed compared to the biofilm conditions (Table S1, FIGURES 4-6 & FIGURE S2). Still, a remarkably number of reads was mapped to the *pet6* gene under all tested conditions (FIGURE 3). In general, the transcription level of *pet6* was low and close to 0.4 % of the transcription observed for *rpoD* (TABLE S2). In summary the data imply that *pet*6 gene expression is not strongly activated by any of the added natural polymers and compared to PET powder (1% w/v) containing cultures. However, the expression was significantly higher in the planktonic cultures compared to the biofilm cultures (FIGURE 3).

### *Pet6* and *ulaG* transcription are induced by BHET in DSM 21264

Since PET6 hydrolyzed PET, we further asked if DSM 21264 would be able to metabolize BHET and use it as sole carbon and energy source, and if it would affect *pet6* gene expression. To address these questions additional growth experiments in liquid media were performed challenging DSM 21264 with BHET at concentrations of 0.5, 5.0 and 30 mM and analyzing the overall gene expression profile using RNAseq. As controls we grew DSM 21264 in the presence of 1.0 mM TPA, DMSO (0.5 % v/v) and PET powder (1 % w/v) in liquid cultures. TABLE S2 summarizes the data obtained for all RNAseq experiments. The volcano plots highlight major changes in gene expression (FIGURE S2). A detailed analysis identified a large number of genes and operons differentially regulated (>log2-foldchange of 2) in the presence of the different BHET concentrations clearly indicating that DSM 21264 senses and adapts its metabolism to the presence of BHET (TABLE S1 and FIGURES 4-6). High concentrations (5.0 and 30 mM) of BHET had the most pronounced effects on the overall gene expression levels in DSM 21264 with >230 upregulated and >100 downregulated genes in the presence of 5.0 mM and >830 upregulated and >550 downregulated genes in the presence of 30 mM BHET compared to the DMSO or the TPA controls (FIGURE S7). Our data imply that BHET is most likely initially metabolized by PET6 releasing MHET as the primary degradation product of BHET. This is in line with an almost 5-fold (log2-foldchange) increased transcription level of *pet6* (AAC977_05355) at 5 mM BHET and 1.8-fold increase at 30 mM. Surprisingly, the RNAseq data indicated that UlaG, a predicted metallo-ß-lactamase, is also involved in BHET metabolism. In the presence of 30 mM BHET *ulaG* was most strongly transcribed with a 7.9-fold upregulation compared to the control. To further verify this novel role of UlaG, we expressed it in *E. coli* T7 SHuffle and used the recombinant protein to hydrolyze BHET. In these tests the promiscuous enzyme was able to cleave BHET at slow but significant rates (FIGURE S3). Further UHPLC measurement confirmed that MHET is the main and initial degradation product of DSM 21264 cells grown on BHET (FIGURE S4). While MHET is converted to EG and TPA, DSM 21264 was not able to grow on TPA and EG as sole carbon and energy source. It does not encode any mono- and dioxygenases involved in aromatic ring cleavage.

### BHET and TPA affect the central circuits involved in c-di-GMP, cAMP-CRP and QS signaling in DSM 21264

During the above-described growth experiments using BHET as a substrate, we noticed major phenotypic changes in DSM 21264 affecting colony morphology, biofilm formation, prodigiosin biosynthesis and others (FIGURE S6).

BHET altered colony morphology already at rather low concentrations (0.5 mM). Colonies were less structured and had smooth edges. Higher concentrations of BHET resulted in the disappearance of the red pigment prodigiosin. Biofilm formation was disturbed at 0.5 mM and no solid biofilms were formed at 10 mM BHET present in the medium (FIGURE S6).

These phenotypic observations implied a larger impact of BHET on the metabolism and main regulatory circuits of DSM 21264 in planktonic cultures. To further elucidate these effects, we analyzed our transcriptome data with respect to the influence of BHET on regulatory pathways linked to QS, cAMP-CRP signaling and c-di-GMP signaling. These pathways were chosen, as they are known to have a wider impact on biofilm formation, motility, secretion, secondary metabolite production and others.

Most notably, 30 mM BHET strongly affected the transcription of genes essential for c-di-GMP biosynthesis. The signal c-di-GMP is synthesized by diguanylate cyclases and DSM 21264 harbors at least 16 possible c-di-GMP cyclases in its genome. The majority of the diguanylate cyclases were strongly downregulated in their transcription in the presence of high concentrations of BHET, while they were strongly upregulated in the presence of low concentrations of BHET or other carbon sources (FIGURE 4). The most strongly attenuated genes coding for diguanylate cyclases were *cdgB, C, I, J* and *M* (log2-foldchange < –4.5). On the contrary, *cdgG* and *acgB* were strongly upregulated in the presence of increased BHET concentrations (log2-foldchange 3.5 and 5) (FIGURE 4).

**FIGURE 4:**
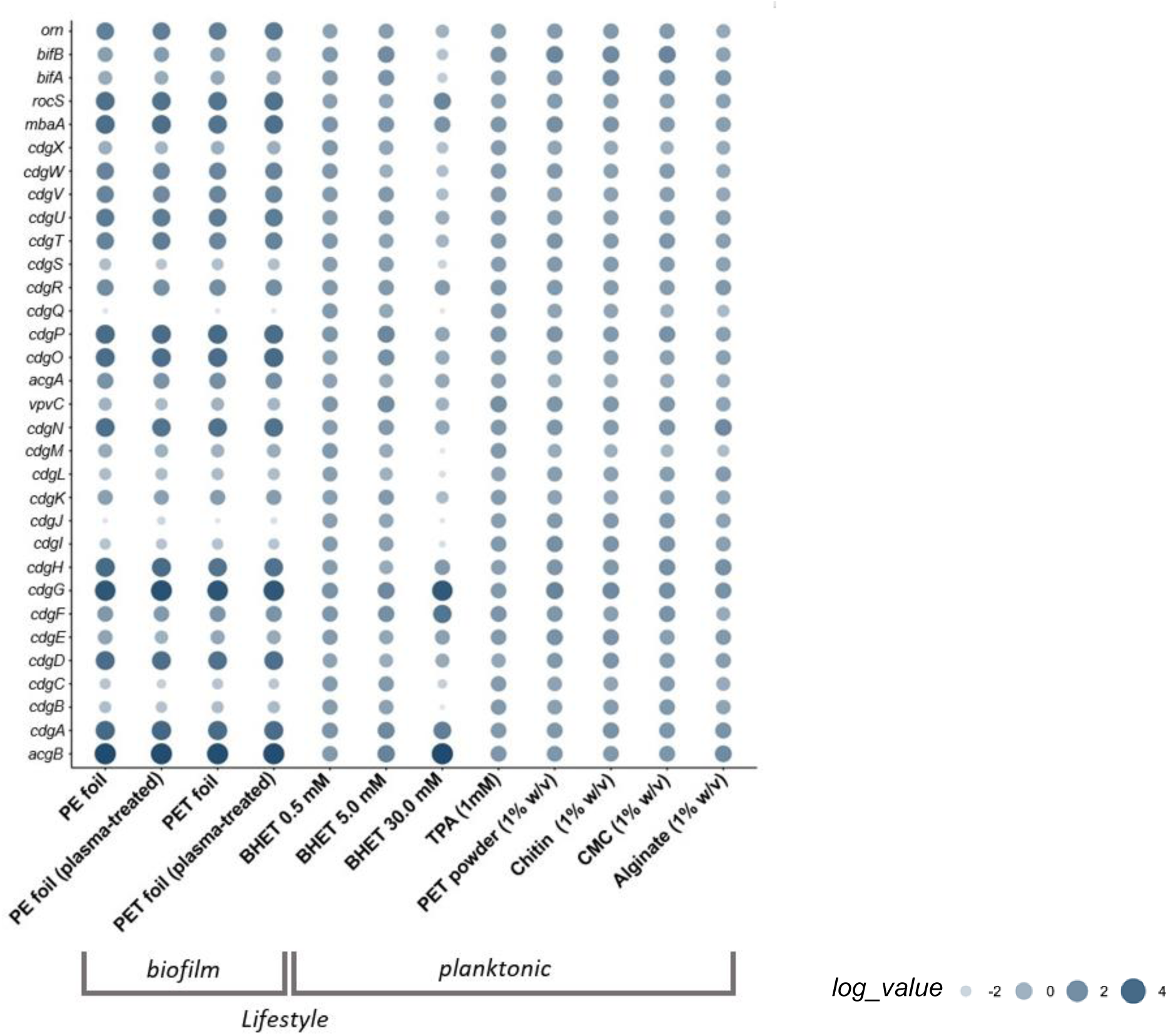
Relative transcription level of c-di-GMP-signaling related genes in DSM 21264. The circle size and color correlate with the normalized transcription level. The color intensity and size of the circles is adjusted to logarithmized values (log_value) ranging from −2 to 4 according to the mapped reads. CdgA-CdgN are diguanylate cyclases (DGC) and CdgO-CdgX are phosphodiesterases (PDE), *bifA* and *bifB* code for bifunctional genes and Orn is an oligoribonuclease. Each data point is a mean value of three independent experiments for each of the 12 conditions shown.

Further, we observed that BHET affected quorum sensing (QS) processes. In this context, transcription of the master QS regulator *aphA* was almost not detectable at 30 mM BHET, while it was well expressed at 0.5 and 5.0 mM BHET or in the presence of TPA in planktonic cultures (FIGURE 5). HapR was also only weakly transcribed in the presence of 30 mM BHET. While it is well known that AphA and HapR are usually transcribed at opposite levels, no differences in transcription levels were visible neither in the presence of lower concentrations of BHET nor in the presence of TPA. These differences in expression were, however, clearly visible in all the biofilm experiments and in the planktonic cultures supplemented with natural polymers (FIGURE 5). Furthermore, TPA negatively affected the transcription of the autoinducer synthase gene *cqsA,* which is involved in the cholera autoinducer I biosynthesis (CAI-I) (TABLE S1 and FIGURE 5).

**FIGURE 5:**
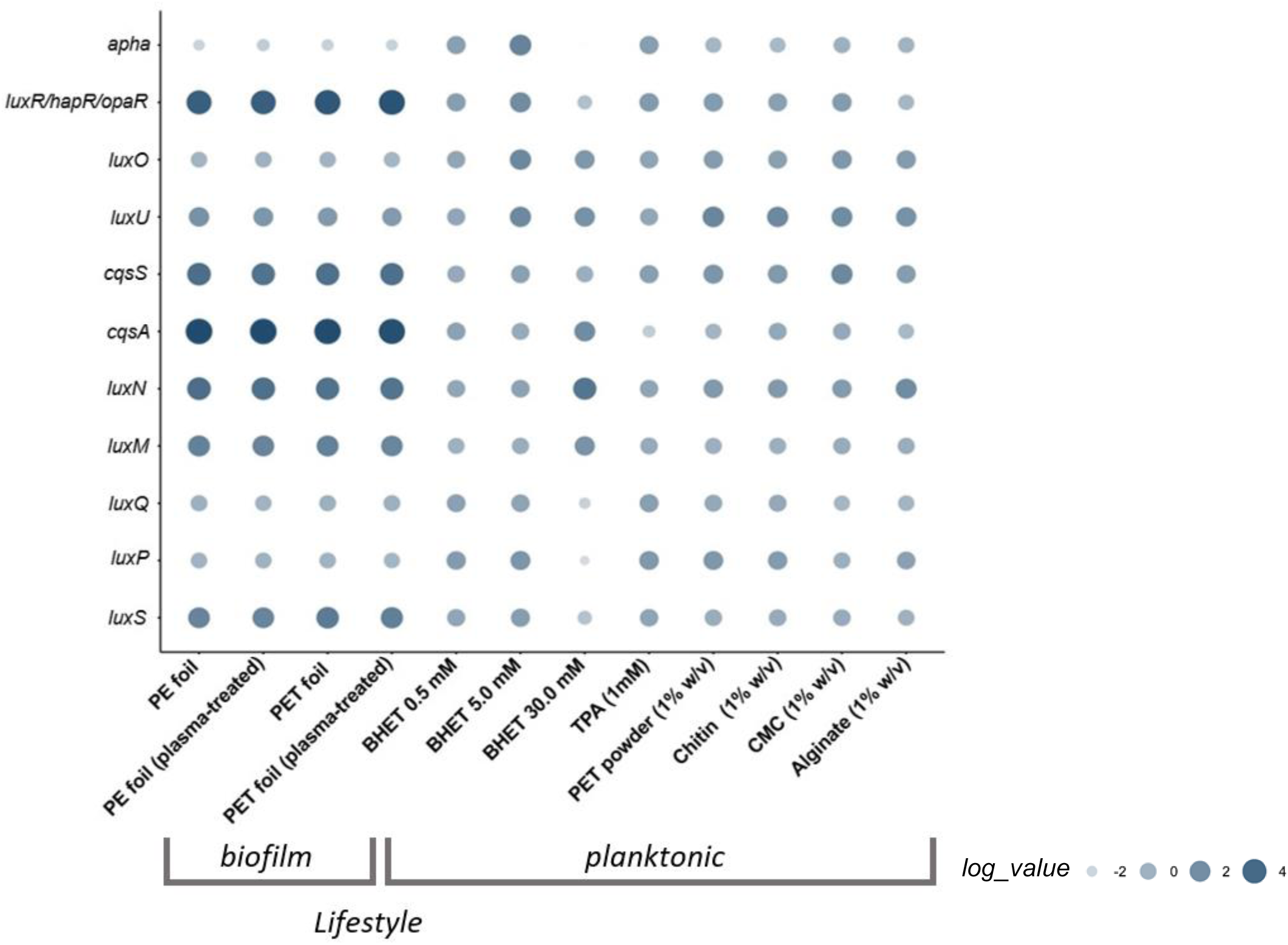
Relative transcription level of essential quorum sensing (QS)-related and differentially regulated genes in DSM 21264. The circle size and color correlate with the normalized transcription level. The color intensity and size of the circles is adjusted to logarithmized values (log_value) ranging from −2 to 4 according to the mapped reads. *luxS*, *luxM* and *cqsA* code for autoinducer synthases, LuxP is annotated as periplasmatic binding protein and LuxQ, LuxN and CqsS are autoinducer sensor kinases. Apha is the low cell density regulator and LuxR high cell density regulator. Each data point is a mean value of three independent experiments for each of the 12 conditions shown.

Further our data showed that the DSM 21264 autoinducer (AI) -I synthase, LuxM, and the cognate sensor LuxN as well as the genes coding for the AI-2 synthase, *luxS*, the periplasmatic binding protein, LuxP, and the sensor kinase LuxQ were significantly affected in their transcription levels at 30 mM BHET (FIGURE 5).

Finally, our data implied that BHET interfered with the cAMP-CRP signaling. The catabolite activator protein transcription was almost completely turned off in the presence of 30 mM BHET (FIGURE 6).

**FIGURE 6:**
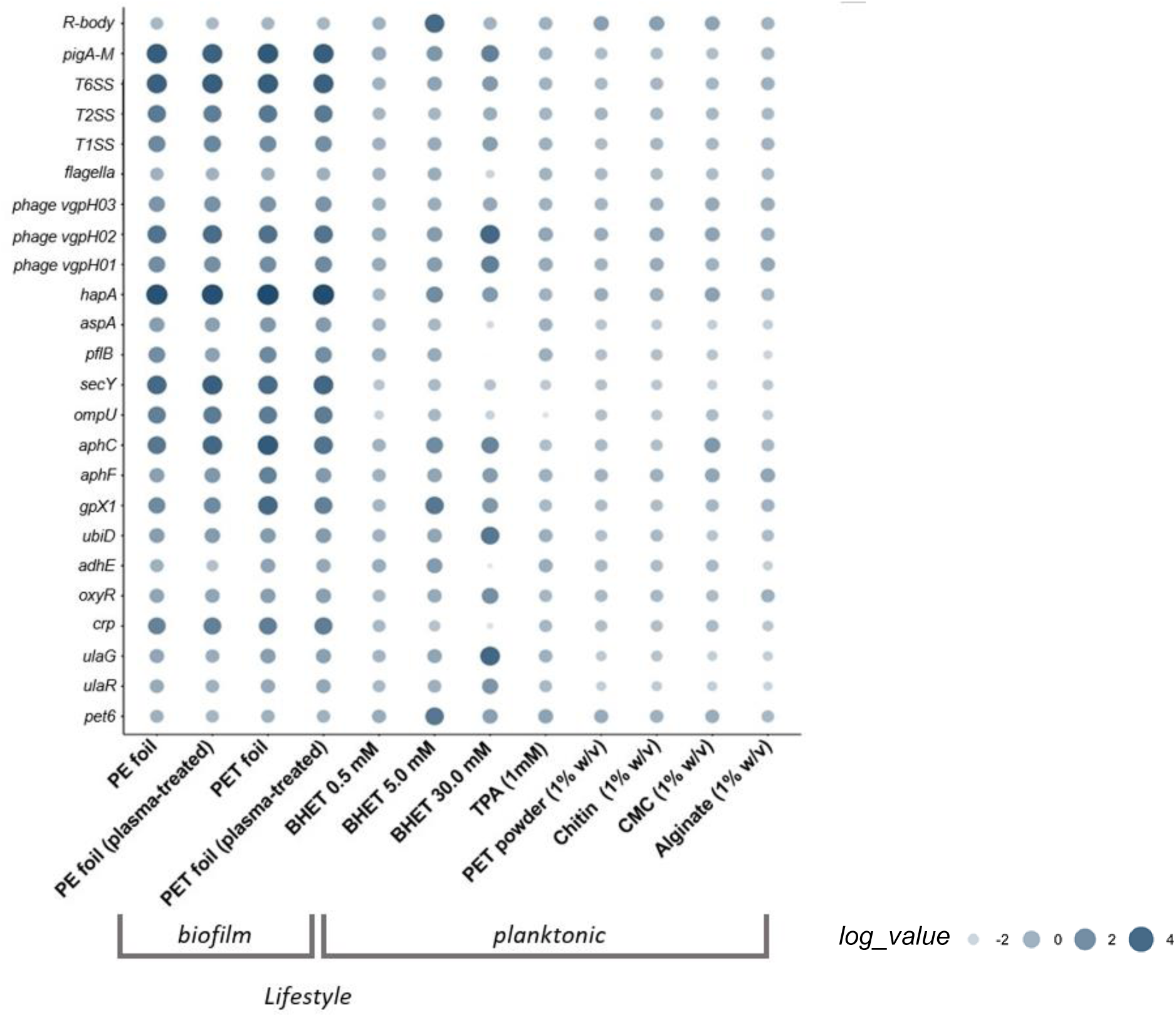
Relative transcription level and changes of differentially regulated major genes and gene clusters in DSM 21264. The circle size and color correlate with the normalized transcription level. The color intensity and size of the circles is adjusted to logarithmized values (log_value) ranging from −2 to 4 according to the mapped reads. R bodies belong to the Rebb family gene cluster, pig gene cluster is responsible for synthesis of prodigiosin, hapA is M4 metallopeptidase annotated as vibriolysin, *aspA* encodes an aspartate ammonia-lyase, *pflB* codes for a formate C-acetyltransferase. Each data point is a mean value of three independent experiments for each of the 12 conditions shown.

Besides the direct impact of 30 mM BHET on the above-mentioned regulatory circuits, 30 mM BHET strongly attenuated transcription of motility genes, *hap*A, *aspE, ompU*. In the contrary, *ulaG, ulaR*, *aphC*, *aphF* and the prodigiosin operon were strongly upregulated (FIGURE 6).

DSM 21264 codes for three prophages (Phage 1, designated VGPH01, AAC977_19445 – AAC977_19700; Phage 2, designated VGPH02, AAC977_20360 – AAC977_20490 and Phage 3, designated VGPH03, AAC977_09080 – AAC977_09210) which can be grouped into the class of Caudoviricetes. VGPH01 and VGPH02 are encoded on the smaller chromosome CP151641 whereas VGPH03 is part of the larger chromosome CP151640. All three phages were transcribed under biofilm conditions. In contrast, their transcription was generally low in planktonic cultures. However, in planktonic cultures, 30 mM BHET resulted in strong transcription of the gene clusters coding for VGPH02 (TABLE S4 and FIGURE 6).

In summary our data imply that BHET and TPA may have a wider effect on the metabolism of DSM 21264 and interfere with the interconnected pathways involved in c-di-GMP, QS and cAMP-CRP signaling.

## DISCUSSION

Today 113 PET-active enzymes are known and most of these enzymes have been characterized very well with respect to their structures, functions and catalytic activities on the synthetic polymer. These enzymes are generally secreted and promiscuous hydrolases belonging to the E.C. 3.1.-. [12] [11] [13] [14] [6]. However, only very few studies have analyzed growth of bacteria on PET foil using global RNAseq approaches [36] [37] [38]. Despite these first studies, it is not clear if bacteria differentiate between the different types of synthetic polymers either as a surface to attach to or as a potential substrate for breakdown.

Our data suggest that only very few genes are differentially regulated when DSM 21264 is grown on PET versus the non-biodegradable PE, implying that most likely *Vibrio gazogenes* is not able to differentiate between these two polymers. Also, the treatment with plasma did not affect this response. Interestingly, our data identified *ompU* as one of the most strongly transcribed genes when grown in biofilms (TABLE 2). OmpU is an outer membrane protein known to be an important adherence factor in *Vibrio cholerae* and an essential biofilm matrix assembly protein [39]. Based on these earlier findings it is likely that the DSM 21264 OmpU plays a major role in biofilm formation on plastic surfaces and life in the plastisphere.

**TABLE 2:**
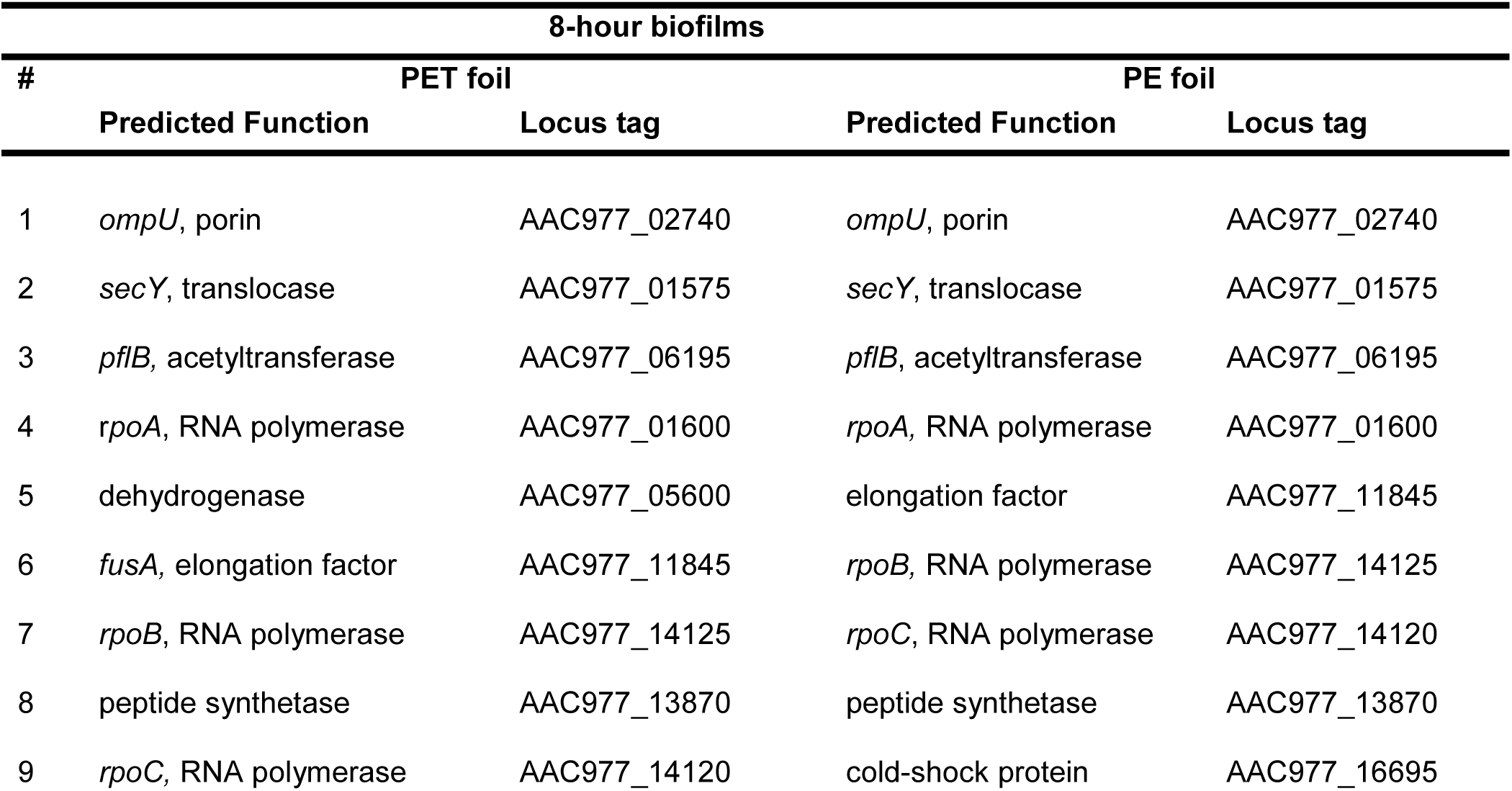
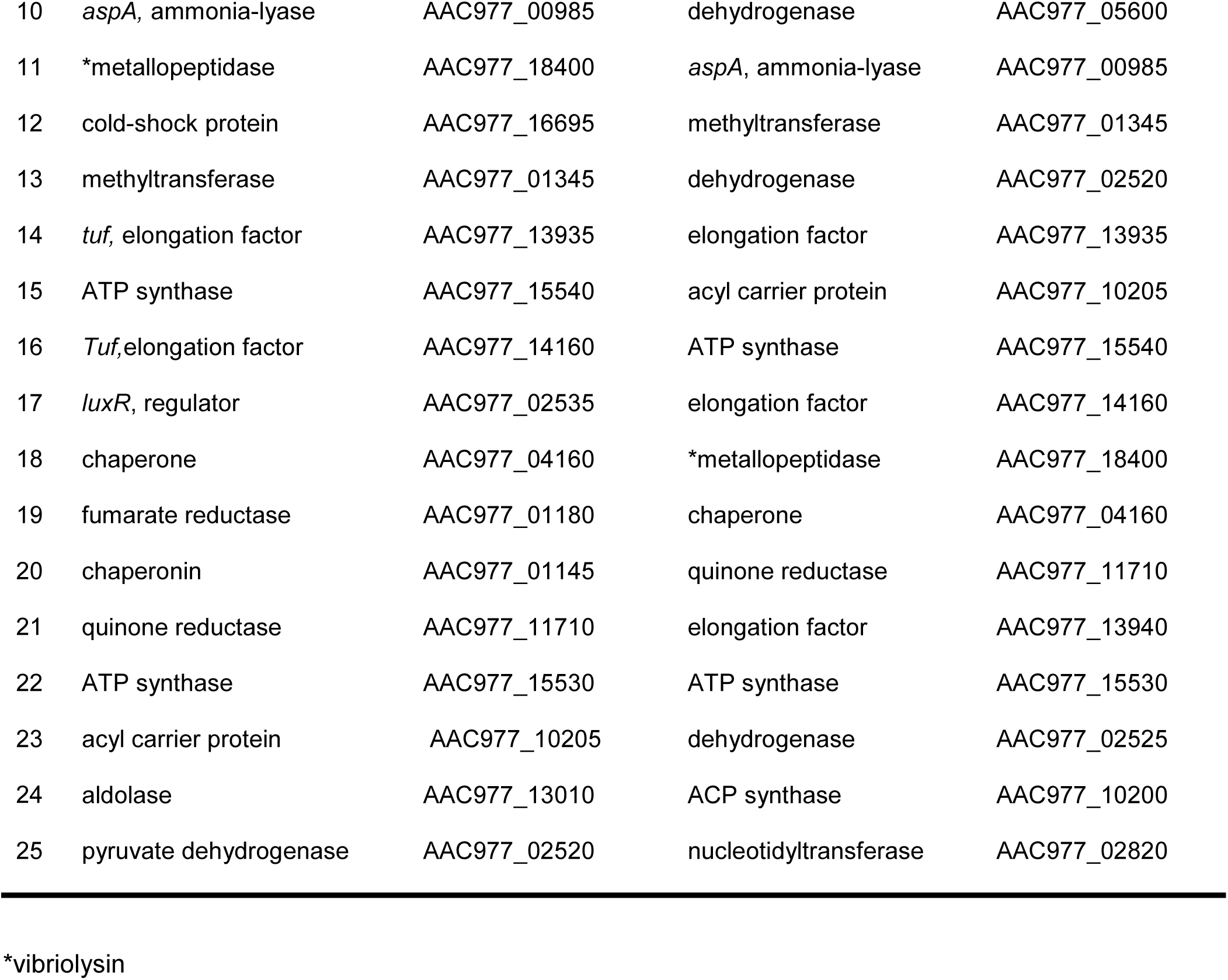
The 25 genes with the highest normalized counts in DSM 21264 grown in biofilms on PET and PE.

| 8-hour biofilms |  |  |  |  |
| --- | --- | --- | --- | --- |
| # | PET foil |  | PE foil |  |
|  | Predicted Function | Locus tag | Predicted Function | Locus tag |
| 1 | *ompU*, porin | AAC977_02740 | *ompU*, porin | AAC977_02740 |
| 2 | *secY*, translocase | AAC977_01575 | *secY*, translocase | AAC977_01575 |
| 3 | *pflB*, acetyltransferase | AAC977_06195 | *pflB*, acetyltransferase | AAC977_06195 |
| 4 | *rpoA*, RNA polymerase | AAC977_01600 | *rpoA*, RNA polymerase | AAC977_01600 |
| 5 | dehydrogenase | AAC977_05600 | elongation factor | AAC977_11845 |
| 6 | *fusA*, elongation factor | AAC977_11845 | *rpoB*, RNA polymerase | AAC977_14125 |
| 7 | *rpoB*, RNA polymerase | AAC977_14125 | *rpoC*, RNA polymerase | AAC977_14120 |
| 8 | peptide synthetase | AAC977_13870 | peptide synthetase | AAC977_13870 |
| 9 | *rpoC*, RNA polymerase | AAC977_14120 | cold-shock protein | AAC977_16695 |
| 10 | *aspA*, ammonia-lyase | AAC977_00985 | dehydrogenase | AAC977_05600 |
| 11 | *metallopeptidase | AAC977_18400 | *aspA*, ammonia-lyase | AAC977_00985 |
| 12 | cold-shock protein | AAC977_16695 | methyltransferase | AAC977_01345 |
| 13 | methyltransferase | AAC977_01345 | dehydrogenase | AAC977_02520 |
| 14 | *tuf*, elongation factor | AAC977_13935 | elongation factor | AAC977_13935 |
| 15 | ATP synthase | AAC977_15540 | acyl carrier protein | AAC977_10205 |
| 16 | *Tuf*, elongation factor | AAC977_14160 | ATP synthase | AAC977_15540 |
| 17 | *luxR*, regulator | AAC977_02535 | elongation factor | AAC977_14160 |
| 18 | chaperone | AAC977_04160 | *metallopeptidase | AAC977_18400 |
| 19 | fumarate reductase | AAC977_01180 | chaperone | AAC977_04160 |
| 20 | chaperonin | AAC977_01145 | quinone reductase | AAC977_11710 |
| 21 | quinone reductase | AAC977_11710 | elongation factor | AAC977_13940 |
| 22 | ATP synthase | AAC977_15530 | ATP synthase | AAC977_15530 |
| 23 | acyl carrier protein | AAC977_10205 | dehydrogenase | AAC977_02525 |
| 24 | aldolase | AAC977_13010 | ACP synthase | AAC977_10200 |
| 25 | pyruvate dehydrogenase | AAC977_02520 | nucleotidyltransferase | AAC977_02820 |

\*vibriolysin

For natural polymers like cellulose, starch and chitin it is well known that the dimers or monomers as primary degradation products are involved in activation of the pathways and transporters essential to metabolize the respective polymers [40] [41] [42]. The activation of the genes involved in the catabolism of the respective monomers is subject to regulation by cAMP-CRP and in part c-di-GMP signaling [43] [44] [45]. Thereby, especially chitin degradation is well studied in *Vibrio* spp. [44] [46]. In contrast little is known about PET or any other synthetic polymer. Recent work implied that the transcriptional regulator MRP (MHETase gene-regulating protein) positively controls the expression of the gram-negative *Idonella sakaiensis* MHETase and the genes involved in TPA metabolism. This first study suggested that MRP possibly mediates the PET-dependent induction of the MHETase and the mono- and dioxygenases genes involved in TPA degradation [47]. These findings are supported by the earlier observation made for *Commamonas thioxidans* strains E6 and S23 [48]. In *C. thiooxidans* the *tphC-A* operon is involved in binding to TPA in the periplasm and its transcription is regulated in the presence of TPA at nM levels [49] [50, 51]. Notably, both strains do not carry active PETase genes. However, they are both able to import TPA and grow on it as a single carbon and energy source [52] [49].

The data from our study suggests that BHET and TPA are nutritional signals affecting QS, c-di-GMP and cAMP signaling at mM concentrations (FIGURE 4-7). This hypothesis is based on 42 transcriptome datasets of DSM 21264 that have given us a very deep insight into the metabolic activities of this bacterium in response to PET, other polymers, BHET and TPA. Because of the important role of *Vibrio* spp. in pathogenicity regulatory networks have been well studied in this genus [53] [54] [27]. Only a few *Vibrio* species harbor PET6 homologues but all of them share a very conserved core genome and have common regulatory networks [31] [55] [56].

Within this setting our data imply that *pet6* is expressed at low levels under most environmental conditions (FIGURE 3) and increasing BHET concentrations affected transcription of the gene. Based on our transcriptome and experimental data we present strong evidence that both BHET and TPA are nutritional signals interfering with the three main regulatory and interconnected circuits involved in QS, cAMP-CRP and c-di-GMP signaling (TABLE S1 and FIGURES 4-6). These regulatory networks are known to be involved in surface attachment, biofilm formation, infection, toxin and secondary metabolite production, phage assembly and others in *Vibrio* spp.. Biofilm formation in *Vibrio* spp. is controlled by an integrated and highly complex network of multiple transcriptional activators (e.g. VpsR, VpsT, and AphA) and transcriptional repressors (HapR, H-NS); but also, regulatory small RNAs and alternative RNA polymerase sigma factors are involved in the regulation of this process. The c-di-GMP signaling influences the planktonic-to-biofilm transition as it positively regulates biofilm formation and negatively regulates motility. In addition, cAMP-CRP-signaling represses biofilm formation in *Vibrio* spp. [57] [54] [58].

In this study, we showed that many of the above-mentioned regulatory circuits and their key regulators in DSM 21264 were affected by BHET and in part by TPA in planktonic cultures (FIGURES 4-6). For instance, AphA and the transcriptional repressor HapR which are both essential components of the QS signaling were strongly affected by high concentrations of BHET. Like AphA and HapR, the autoinducer-I and 2 synthase genes, the cognate sensor genes and the transcription of the periplasmic binding protein were also significantly affected in their transcription levels by 30 mM BHET (FIGURE 5).

The major signal c-di-GMP is produced by diguanylate cyclases and degraded by specific phosphodiesterases [59] [58]. DSM 21264 has a very similar set of c-di-GMP cyclases and phosphodiesterases compared to *V. cholerae* or other well studied *Vibrio* species. Many of the cyclase genes were attenuated by the higher concentrations of BHET (TABLE S1 and FIGURE 4). This observation is in line with the failure to form biofilms at higher BHET concentrations (FIGURE S6). The lack of sufficient cyclase transcription will most likely result in low intracellular levels of c-di-GMP that would not be sufficient to allow biofilm formation.

Within this framework, cAMP-CRP signaling is well known to be involved in the expression of inducible catabolic operons. Besides this function cAMP-CRP signaling is also involved in processes related to infection and biofilm formation. Thereby, a specific adenylate cyclase catalyzes the formation of cAMP from ATP, whereas cAMP phosphodiesterases catalyze its breakdown [60] [61][54]. The DSM 21264 cAMP-CRP signaling pathway is conserved as well and its main regulatory protein is the CRP regulator. While CRP was highly transcribed under biofilm conditions, its transcription was in general much lower in planktonic lifestyle. Notably, 5.0 and 30 mM BHET downregulated the CRP transcription even further, implying that BHET is interfering with this signaling cycle as well (FIGURE 6).

Besides these observations our study has further revealed that UlaG is involved in BHET metabolism. This novel function of UlaG has previously not been described and while we initially observed a strongly increased transcription of *ulaG* in the presence of high BHET concentrations, we provided experimental data to confirm this novel functional role of UlaG (FIGURE S3). Previous work has already demonstrated that UlaG is a promiscuous metallo-beta-lactamase with a wide range of functions in bacteria and archaea [62] [63]. In addition, our data support the notion that *adhE* is involved in the metabolism of EG. This was confirmed by both RNAseq and experimental data (FIGURE S5). Since *adhE* is transcribed at higher levels in PET biofilms it is likely that this increase is caused already by the degradation of the polymer.

Based on the above made observations we were able to establish a putative first model outlining the main genes and enzymes involved in PET uptake and metabolism and the regulatory circuits involved (FIGURE 7). This preliminary model and our experimental data provide a foundation for future research on bacterial metabolism and nutritional signaling during life in the plastisphere. Since *Vibrio* spp. is found with high frequencies in the plastisphere data from this study will help to identify key factors involved in enriching for pathogens in this man-made habitat. Finally, this study will be useful for the design of bacterial strains for efficient plastics removal in the marine environment.

**FIGURE 7:**
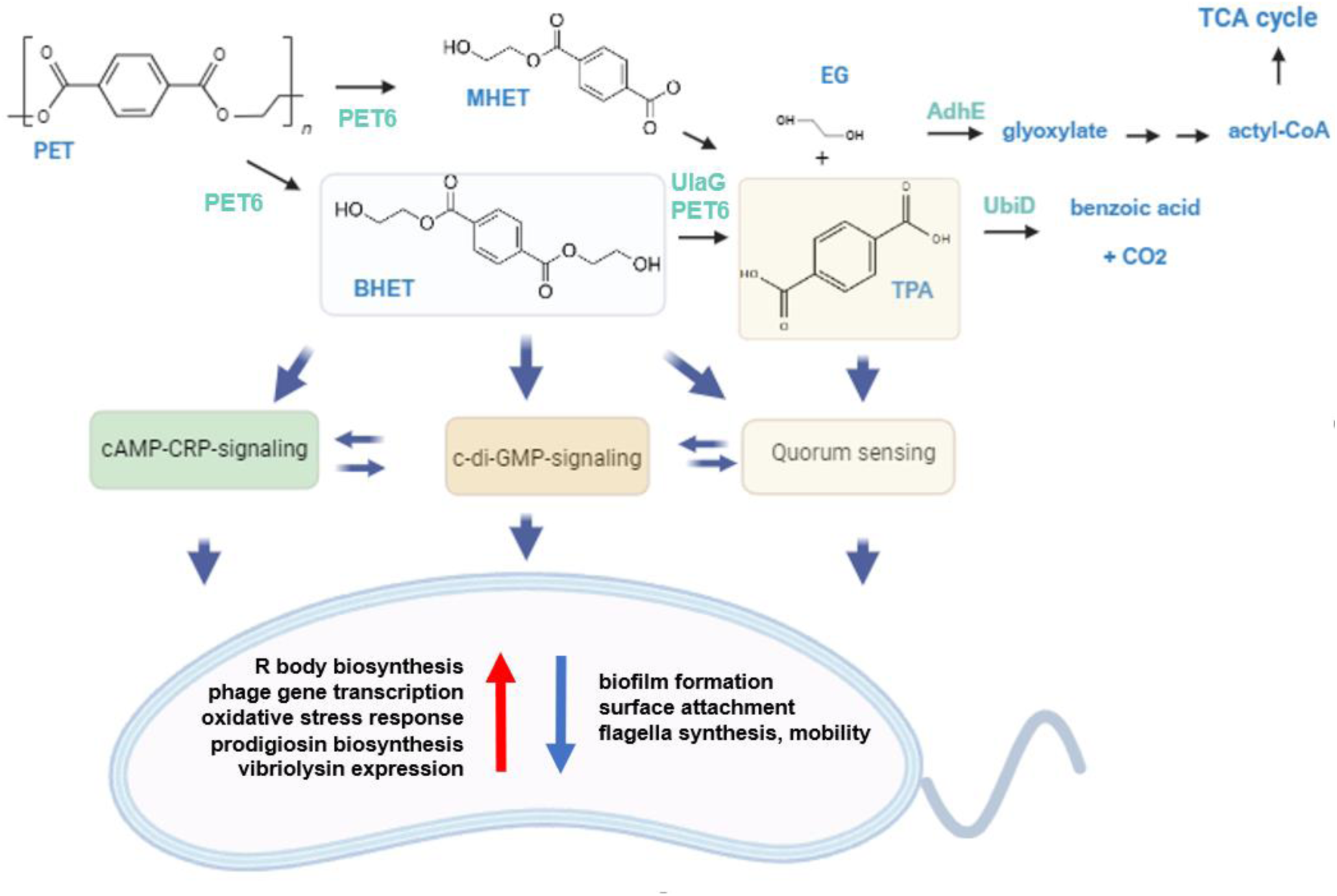
Possible PET degradation pathway in DSM 21264 and regulatory model affecting QS, cAMP-CRP and c-di-GMP signaling through BHET and TPA. Proteins involved in enzymatic PET breakdown are in cyan color. PET and degradation products released are labeled in blue. Blue arrows indicate regulatory pathways affected by BHET and /or TPA in DSM 21264. Red and blue arrows inside the cell model indicate up- and downregulated pathways.

## ACKNOWLEDGEMENTS

This work was funded in part by the University Hamburg and in supported by the European Commission (Horizon2020 project FuturEnzyme; grant agreement ID 101000327).

L. Preuß, C. Vollstedt and W.R. Streit conceived the study. L. Preuß, A. Dumnitch, Ly Trinh, C. Vollstedt and W.R. Streit generated the data. W.R. Streit acquired funding. M. Alawi, A. Poehlein and R. Daniel carried out the bioinformatic analyses. N. Burmeister and W. Maison provided the plasma-treated surfaces.

## DATA AVAILABILITY

Sequence data reported in this publication have been submitted to NCBI/ENA. The raw reads of the 42 sequencing runs are publicly available under accession PRJEB80907 at European Nucleotide Archive (ENA). The newly established genome sequence of *Vibrio gazogenes* DSM 21264 is available under accession numbers CP151640 and CP151641 at NCBI/GenBank.

## MATERIAL AND METHODS

### Bacterial strains and growth conditions

Bacterial strains and plasmids used in this study are summarized in TABLE 1. *Vibrio* sp. was cultured either at 22 °C or 28 °C in artificial seawater medium (28.13 g/L NaCl, 0.77 g/L KCl, 1.6 g/L CaCl_2_ x 2H_2_O, 4.8 g/L MgCl_2_ x 6 H_2_O, 0.11 g/L NaHCO_3,_ 3.5 g/L MgSO_4_ x 7 H_2_O, 10 g/L yeast extract, 10 g/L tryptone) or in AS medium 1:10 diluted (1 g/L tryptone, 1 g/L yeast extract) and either CMC (1 % w/v), chitin (1 % w/v), alginate (1 % w/v), PET powder (1 % w/v), TPA (1mM) or BHET added (0.5, 5 and 30 mM). Additional growth experiments were implemented in M9 medium (Na_2_HPO_4_ 33.7 mM, KH_2_PO_2_ 22 mM, NaCl 51.3 mM, NH_4_Cl 9.35 mM, MgSO_4_ 1 mM, biotin 1 µg, thiamin 1 µg, EDTA 0.134 mM, FeCl_3_-6H_2_O 0.031 mM, ZnCl_2_ 0.0062 mM, CuCl_2_ 2H_2_O 0.76 µM, CoCl_2_ 2 H_2_O 0.42 µM, H_3_BO_3_ 1.62 µM, MnCl_2_ 4H_2_O 0.081 µM).

*E. coli* was grown at 37°C aerobically in LB medium (10 g/L tryptone, 5 g/L yeast, 5 g/L NaCl) and supplemented with the appropriate antibiotics.

For the observation of the phenotypical reaction and halo formation of DSM 21264 to various concentrations of BHET LB agar plates (10 g/L tryptone, 5 g/L yeast, 5 g/L NaCL, 15 g/L agar) were prepared. After autoclaving the respective amount of BHET was added to reach final concentrations of 0.5 mM, 5.0 mM, 10 mM and 30 mM in the respective plate. 5 µl of DSM 21264 OD_600 nm_ = 1 were added and incubated 24 h at 28 °C.

### Fluorescence imaging analysis of biofilms

For observation of biofilm formation of DSM 21264 on plastics surfaces cells were inoculated at OD_600 nm_ = 0.05 in ASW medium and incubated in 6 -well cell culture plates (Nunc cell culture plate, catalog no. 130184; Thermo Fisher Scientific, Waltham, MA) at 22 °C or 28 °C at 80 rpm shaking. After incubation foil was washed in PBS buffer and placed into µ-slide eight-well (ibiTreat. catalog no. 80826, ibidi USA, Inc., Fitchburg, Wisconsin). Cells were stained using LIVE/DEAD BacLight bacterial viability kit (Thermo Fisher Scientific, Waltham, MA, USA).

Cells were visualized using confocal laser scanning microscope (CLSM) Axio Observer.Z1/7 LSM 800 with airyscan (Carl Zeiss Microscopy GmbH, Jena, Germany) and a C-Apochromat 63x/1.20W Korr UV VisIR objective. Settings for the microscope are presented in Table S5. For the analysis of the CLSM images the ZEN software was used (version 2.3. Carl Zeiss Microscopy GmbH, Jena, Germany). For each sample at least three different positions were observed and one representative CLSM image was chosen.

### Preparation for scanning electron microscopy

Strains and polymers were incubated as described above and after incubation fixed overnight in 1 % PFA in 50 mM Cacodylate buffer, pH 7. The following day samples were incubated in 0.25 % GA in 50 mM Cacodylate buffer, pH 7 overnight. For drainage samples were incubated in 30 %, 50 % and 70 % of ethanol in the following order each for 20 min. An additional incubation in 70 % ethanol was performed overnight. Drainage continued with sample incubation in 80 %, 96 % and 100 % of ethanol for 20 min and 100 % of ethanol for 30 min. Critical point drying was performed using Leica EM CPD300 thereby washed 18 x with CO_2._ Next, coated in a thin carbon layer using Sputter Coater LEICA EM ACE600. Microscopy was performed at the scanning electron microscope LEO 1525 using Software SmartSEM V06.00.

### Sample preparation for RNA seq and analysis

Precultures of DSM 21264 were inoculated at OD_600 nm_ = 0.05 in ASW medium (4 mL/well) in 6 - well cell culture plates (Nunc cell culture plate, catalog no. 130184; Thermo Fisher Scientific, Waltham, MA). To each well either PE or PET foil (35 mm diameter) was added and incubated for 8 hours at 22 °C shaking at 80 rpm. Foil was washed in PBS buffer to get rid of planktonic cells, then cells were scratched off the surface. 2 ml of 20 % stop mix consisting of 95 % ethanol and 5 % phenol was added to the cells and the mixture was centrifuged for 20 minutes at 4 °C. Supernatant was discarded, the pellet was washed three times in PBS buffer and afterwards immediately frozen in liquid nitrogen and stored at −70 °C.

The liquid cultures were inoculated at OD_600 nm_ = 0.05 in diluted artificial seawater. Respective C-source (CMC, PET powder (particle size 300 µm max; >50 % crystallinity, product no. ES30-PD-000132), chitin and alginate (1% w/v) and BHET concentrations ranging from 0.5 mM, 5 mM and 30 mM as well as 1 mM TPA) was added and incubated shaking at 28 °C at 130 rpm for 24 hours. As negative controls no additional carbon source and DMSO controls were prepared. Cells were harvested and cell pelleted.

Harvested cells were re-suspended in 800μl RLT buffer (RNeasy Mini Kit, Qiagen) with β-mercaptoethanol (10μl ml^−1^) and cell lysis was performed using a laboratory ball mill. Subsequently, 400μl RLT buffer (RNeasy Mini Kit Qiagen) with β-mercaptoethanol (10μl ml^−1^) and 1200 μl 96 % (vol./vol.) ethanol were added. For RNA isolation, the RNeasy Mini Kit (Qiagen) was used as recommended by the manufacturer, but instead of RW1 buffer RWT buffer (Qiagen) was used in order to isolate RNAs smaller than 200 nucleotides also. To determine the RNA integrity number the isolated RNA was run on an Agilent Bioanalyzer 2100 using an Agilent RNA 6000 Nano Kit as recommended by the manufacturer (Agilent Technologies, Waldbronn, Germany). Remaining genomic DNA was removed by digesting with TURBO DNase (Invitrogen, Thermo Fischer Scientific, Paisley, UK). The Illumina Ribo-Zero plus rRNA Depletion Kit ((Illumina Inc., San Diego, CA, USA) was used to reduce the amount of rRNA-derived sequences. For sequencing, the strand-specific cDNA libraries were constructed with a NEB Next Ultra II Directional RNA library preparation kit for Illumina and the NEB Next Multiplex Oligos for Illumina (96) (New England BioLabs, Frankfurt am Main, Germany). To assess quality and size of the libraries samples were run on an Agilent Bioanalyzer 2100 using an Agilent High Sensitivity DNA Kit as recommended by the manufacturer (Agilent Technologies). Concentration of the libraries was determined using the Qubit® dsDNA HS Assay Kit as recommended by the manufacturer (Life Technologies GmbH, Darmstadt, Germany). Sequencing was performed on the NovaSeq 6000 instrument (Illumina Inc., San Diego, CA, USA) using NovaSeq 6000 SP Reagent Kit (100 cycles) and the NovaSeq XP 2-Lane Kit v1.5 for sequencing in the paired-end mode and running 2x 61 cycles. For quality filtering and removing of remaining adaptor sequences, Trimmomatic-0.39 (Bolger et al., 2014) and a cutoff phred-33 score of 15 were used. The mapping against the reference genome of *V. gazogenes* DSM 21264^T^ was performed with Salmon (v 1.10.2) [64]. As mapping back-bone a file that contains all annotated transcripts excluding rRNA genes and the whole genome of the reference as decoy was prepared with a k-mer size of 11. Decoy-aware mapping was done in selective-alignment mode with ‘–mimicBT2’, ‘– disableChainingHeuristic’and‘–recoverOrphans’flags as well as sequence and position bias correction and 10 000 bootstraps. For–fldMean and–fldSD, values of 325 and 25 were used respectively. The quant.sf files produced by Salmon were subsequently loaded into R (v 4.3.2) [65] using the tximport package (v 1.28.0) [66]. DeSeq2 (v 1.40.1) was used for normalization of the reads and fold change shrinkages were also calculated with DeSeq2 [67] and the apeglm package (v 1.22.0) [68]. Genes with a log2-foldchange of +2/−2 and ap-adjust value <0.05 were considered differentially expressed.

Fastp [69] was used to remove sequences originating from sequencing adapters and sequences of low-quality sequences. Reads were then aligned to the *V. gazogenes* DSM 21264 reference assembly [70] using BWA mem. Differential expression analysis was carried out with DESeq2 [67]. A gene was considered significantly differentially expressed in a comparison if the corresponding false discovery rate (FDR) was smaller or equal to 0.05 and the absolute log2-foldchange (|log2FC|) was larger than 2. All software was used with standard parameters. Sequence data reported in this publication have been submitted to the European Nucleotide Archive (ENA). They are publicly available under accession PRJEB80907.

### Cloning and expression of *V. gazogenes* UlaG

*UlaG* (NCBI Ref. Seq. WP_072962133.1) was cloned into pET21a(+) vector (Novagen/Merck) using restriction sites *Nde*I and *Xho*I in front of C-terminal 6x HisTag. Primers were designed using SnapGene (GSL Biotech LLC, San Diego CA, United States) and PCR was performed using parameters indicated by design tool. After PCR cleanup, 7 µl DNA (≈30 ng/μl) were mixed with 1 μl of T4 Ligase Buffer (10x), 1 µl of T4 Ligase (400 U/μl) and 1 µl of purified pET21a(+) plasmid and incubated over night at 6 °C. Ligation construct was transformed into competent *E. coli* T7 Shuffle cells and positive clones were identified were via DNA sequencing (Microsynth seqlab).

### Protein production and purification

For protein production cells were overexpressed by growing inoculated cultures with T7 SHuffle harboring pET21a(+)::*ulaG* or BL21(DE3) harboring pET28a(+)::*pet6* at 37 °C to OD_600nm_ = 0.8. Cells were induced with IPTG to final concentrations of 0.4 mM and incubated at 22 °C over night until cells were harvested by centrifugation at 5.000 g.

For protein purification cell pellet was resuspended in lysis buffer (50 mM NaH_2_PO_4_, 300 mM NaCl, 10 mM imidazole, pH 8.0) and sonicated for cell disruption with Ultrasonic Processor UP200S by *Hielscher* with amplitude 70 % and cycle 0.5. Afterwards, the proteins harboring a sixfold C-terminal histidine tag were purified with nickel-ion affinity chromatography using Ni-NTA agarose (Qiagen, Hilden, Germany). The elution buffer was exchanged against 0.1 mM potassium phosphate buffer pH 8.0 in a 30 kDa Amicon Tube (GE Health Care, Solingen, Germany).

### Measurement of PET and BHET degradation

Precultures of DSM 21264 were inoculated at OD_600 nm_ = 0.05 in 20 mL M9 medium supplemented with 5 mM of glucose and additional 5 mM of BHET. Flasks were incubated shaking at 28 °C and 130 rpm for six days. Each day 1 mL of each sample was taken and measured at the UHPLC. As negative control medium was incubated under same conditions with *E. coli* Dh5α and samples were taken at same timepoints.

For detection of PET powder and foil degradation DSM 21264 was incubated as described above in 6 well plates and also 0.5 mL of sample was taken each day and prepared for UHPLC measurements.

For enzymatic BHET degradation purified enzyme in ranging concentrations was incubated with 300 µM of BHET for 24 h and samples were prepared for UHPLC.

Respective supernatants were prepared and measured at the UHPLC following protocols described previously [71] [49].

### Cloning promoter fusion of *adhE*

Promoter of *adhE* (AAC977_10260) was identified using the software tool published at softberry.com ([BPROM - Prediction of bacterial promoters](http://www.softberry.com/berry.phtml?topic=bprom&group=programs&subgroup=gfindb)). The region 365 bp downstream from the respective gene was chosen and cloned into pBBR1-MCS-1 vector [72] carrying amCyan in the multiple cloning site using restriction enzymes *Sac*I and *Xba*I.

Primer design, PCR, ligation and transformation into competent *E. coli* DH5α cells were performed as described above. Positive constructs were transformed into competent *E. coli* WM3064 cells. Since electroporation didn’t result in sufficient colony amount, the plasmid was conjugated via biparental conjugation into DSM 21264. Cells were grown in overnight cultures. 1 mL of donor strain (*E. coli* WM3064 carrying the respective plasmid) and receptor strain (DSM 21264) were mixed at ratio 1:1 and centrifuged at 5,000 rpm for 8 min. Cells were washed three times in LB medium, resuspended after final centrifugation step in 150 µl of LB medium and spot was given onto LB agar plate supplemented with 300 µM DAP. The plate was incubated overnight at 28 °C and spot was washed off. Dilutions ranging from 1:1 to 1:5 were prepared and plated onto ASWM agar plates supplemented with 25 µg chloramphenicol and 25 µg kanamycin. Colonies were detectable after 2 to 3 days of incubation and inoculated in liquid medium.

### Promoter fusion measurements

For the identification of the influence of ethylene glycol on the expression of *adhE,* DSM 21264 carrying promoter fusion construct pBBR1MCS-1::*adhE::amcyan* as well as wild type strain were inoculated at OD_600 nm_ = 0.05 in M9 medium with 5 mM glucose. Cells were incubated in 48 well-plates (1 mL/well) (Nunc cell culture plate, catalog no. 150687; Thermo Fisher Scientific, Waltham, MA, USA) for 48 h at 130 rpm shaking. For the detection of the influence of ethylene glycol 5 mM were added and strains were grown in the presence with and without EG. After incubation optical density and fluorescence of cyan (450;496) was measured at Synergy HT plate reader using Gen5 software (Biotek, Winooski, USA). Average was evaluated and fluorescence was divided by optical density to detect fluorescence units.

### Plasma activation of PE and PET foil

For plasma activation, an atmospheric air plasma system from Plasmatreat GmbH (Steinhagen, Germany) was used. The atmospheric-pressure plasma was produced by a generator FG5001 with an applied working frequency of 21 kHz, generating a non-equilibrium discharge in a rotating jet nozzle RD1004 in combination with the stainless-steel tip No. 22826 for an expanded treatment width of approximately 22 mm. Additionally, the jet nozzle was connected to a Janome desktop robot type 2300N for repetitious accuracy regarding treatment conditions. The process gas was dry and oil-free air at an input pressure of 5 bar in all experiments.

