## Supplemental Figures S1-S7, Supplemental Tables S2-S5 for "Polyethylene terephthalate (PET) primary degradation products affect c-di-GMP-, cAMP-signaling and quorum sensing (QS) in *Vibrio gazogenes* DSM 21264"

### for

The authors declare no conflict of interest.

**Correspondence**

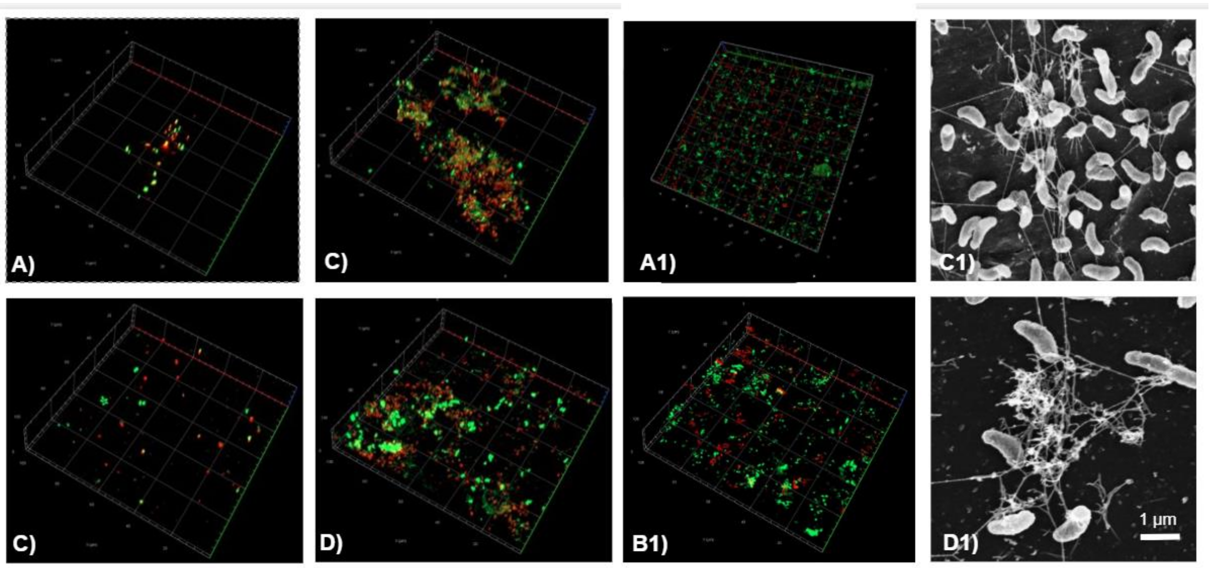


**FIGURE S1:** Confocal laser scanning microscope (CLSM) images of *V. gazogenes* DSM 21246 grown on PET and PE. CLSM images show DSM 21246 grown on PET **A)**, **C)** and PE **B)**, **D)** in ASW salt solution with additional trace elements after 10 days **A)**; **B)** and 180 days **C)**; **D)** of incubation at 22 °C. Figures **A1)**-**D1)** show PE and PET foil previously plasma activated. **A1)** and **B1)** show PE foil, biofilm is evaluated using CLSM and SEM, while **C1)** and **D1)** is DSM 21264 incubated on PET foil.

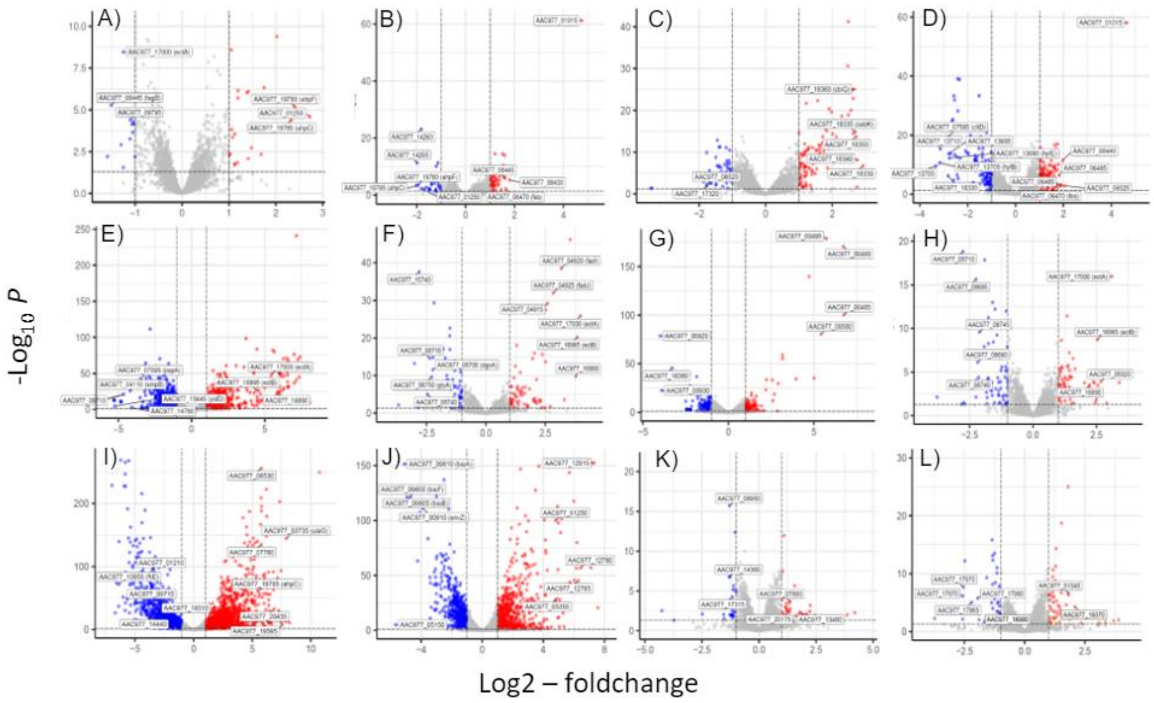

**FIGURE S2:** Volcano plots of *V. gazogenes* transcriptomes and cells grown under different conditions and surfaces. X-axis show log2-foldchange, y-axis show false discovery rate, meaning the significance of the transcripts. Blue dots indicate significantly downregulated genes and red dots significantly upregulated genes. Grey dots indicate not significantly differentially expressed genes. **A)-D)** cells grown in biofilm on plastic surfaces **A)** PET foil vs. PE foil, **B)** PETplasma_PET, **C)** PETplasma_PEplasma, **D)** PEplasma_PE; **E)-L)** transcriptomes of liquid cultures grown with different carbon sources grown in batch cultures in artificial seawater medium at 28°C und under constant shaking at 130 rpM. Biofilms cultures were cultivated as described in 9 tested **E)** alginate**, F)** chitin**, G)** CMC and **H)** PET powder; **I),** 30 mM BHET; **J),** 5 mM BHET; **K),** 0.5 mM BHET and **L)** 1mM TPA. The mRNAs were extracted after 8 hours of growth. The data represent mean values of three individual biological replicates for each carbon source or surface. Locus/ORF tags of most strongly differentiated genes are indicated in the various plots.

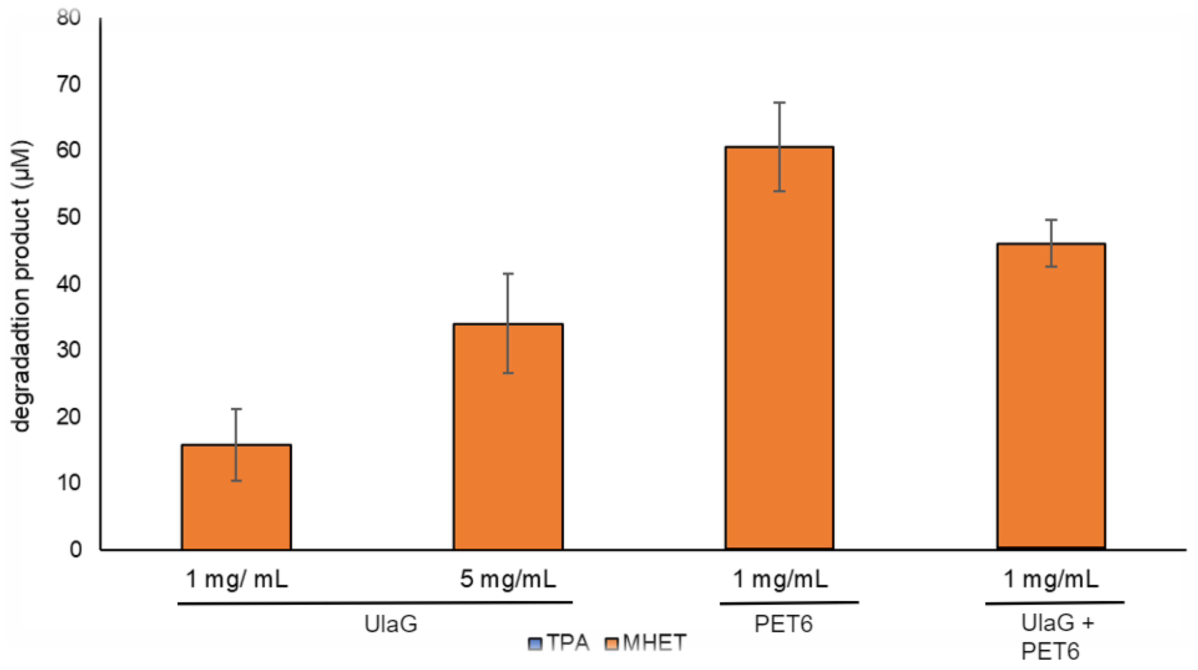


**FIGURE S3**: BHET degradation assay of recombinant and purified UlaG and PET6. Samples were measured using UHPLC ad as outlined in the Materials and Methods section. For detection of degradation products MHET and TPA. BHET was added at 30 mM. Bars indicate production of MHET. Data are mean values of three measurements and the simple SD is given. Data were recorded after 24 hours of incubation.

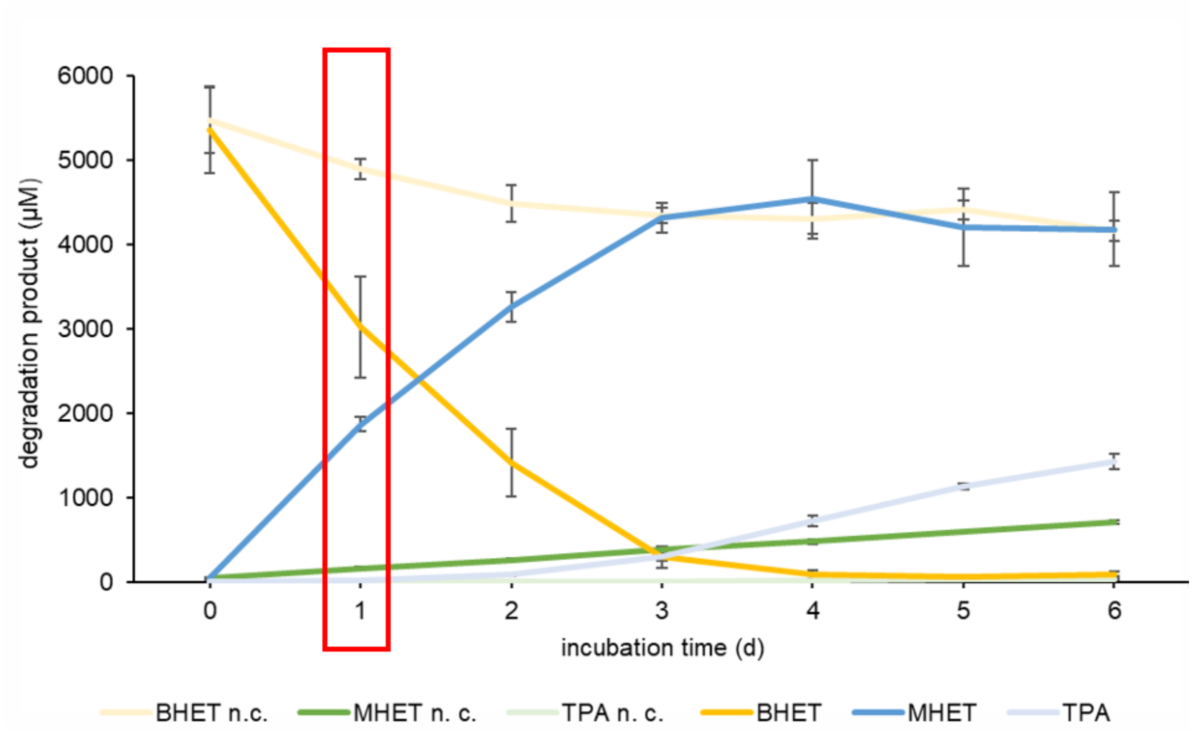


**FIGURE S4:** DSM 21264 incubated with 5 mM BHET in artificial seawater salt solution and incubated with 5 mM of glucose for 6 days at 28 °C at 130 rpm shaking. Graphic shows BHET degradation and its degradation products over time. The red box indicated the timepoint when RNAseq samples were taken. After 3 days BHET is completely degraded into MHET and TPA concentration is increasing. The negative control was *E. coli* DH5α grown at the same OD and under the same conditions.

Data are mean values of five independent measurements and error bars indicate the simple standard deviation.

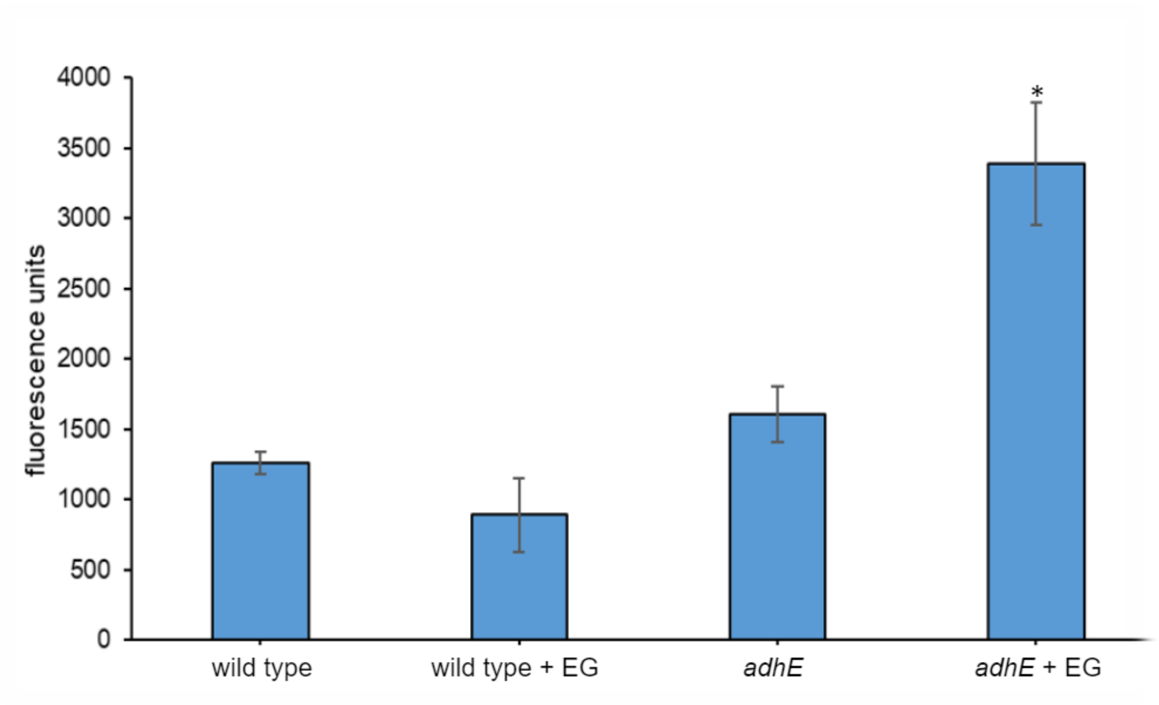


**FIGURE S5:** Incubation of DSM 21264 carrying pBBR1-MCS-1::*adhE*::amcyan in M9 medium (30 g/L NaCl) with 5 mM glucose for 24 h shaking. The promoter fusion of the alcohol dehydrogenase gene *adhE* was induced with 5 mM ethylene glycol. As controls wild type with and without ethylene glycol as well as the promoter fusion without ethylene glycol were used. The promoter fusion constructs induced with ethylene glycol had significantly higher fluorescence values than all used controls at $P<0.01$ and are marked with an asterisk.

Data are mean values of four independent measurements and error bars indicate the simple standard deviation.

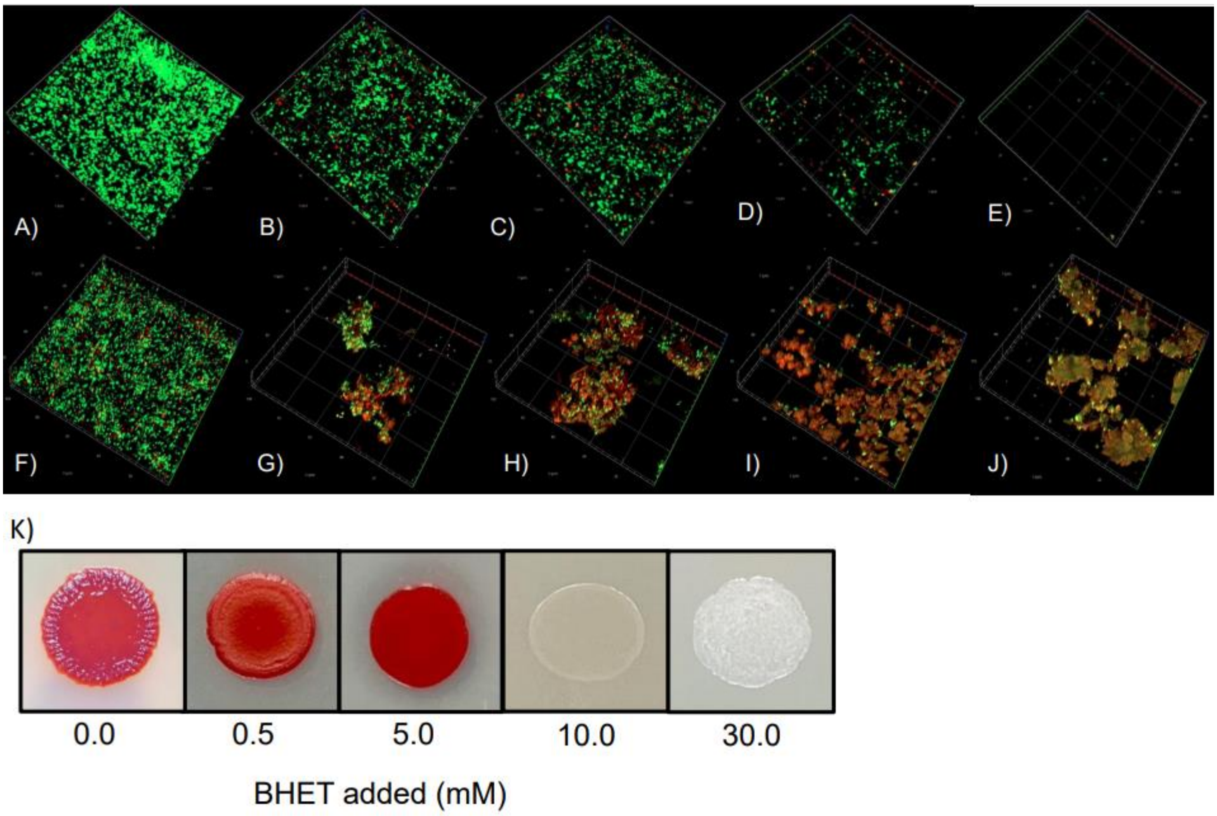


**FIGURE S6:** Confocal laser scanning microscope (CLSM) images **(A-J)** of V. gazogenes DSM 21246 biofilms grown in the presence of increasing concentrations of BHET **(B-E)** and TPA **(G-J)** for 24 h of incubation at 28 °C. BHET concentrations ranged from 0.5 mM **(B)**, 5 mM **(C)**, 10 mM **(D)**, 30 mM **(E)**. Used TPA concentrations were 1 mM **(G)**, 5 mM **(H)**, 10 mM **(I)** and 20 mM **(J)**. As controls DSM 21264 was inoculated with DMSO **(F)** and also without any added substrates **(A)**. Cells were stained using LIVE/DEAD stain.

**K)** Changing colony morphology of DSM 21246 in the presence of increasing BHET concentrations in LB agar plates at 28 °C after 24 h of incubation.

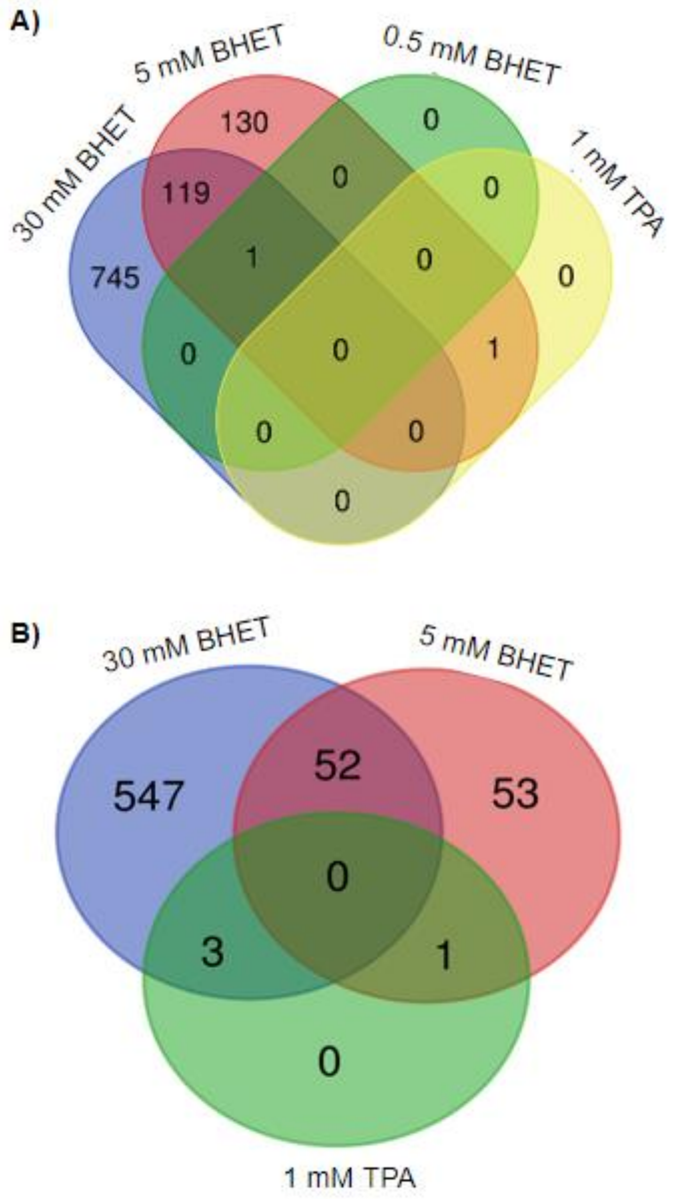


**FIGURE S7**: VENN diagrams of differentially transcribed genes with log2-foldchange above 2. **A)** shows upregulated genes and **B)** downregulated genes regarding the respective condition. Files were evaluated using https://bioinformatics.psb.ugent.be/webtools/Venn/ website.

**TABLE S2: *V. gazogenes* DSM 241264 transcriptomes established and analyzed in this study.**

| Lifestyle | **Average read number** (mio reads) | **Reads mapped** (%) | **Reads mapped to *rpoD*** (*# reads*) | **Reads mapped to *pet6*** (*# reads*)(% of *rpoD*) |
|---|---|---|---|---|
| *Biofilm* | | | | |
| PET foil | 11.0 | 96.3 | 11.879 | 50.0 (0.4) |
| PE foil | 10.1 | 95.9 | 11.772 | 50.0 (0.4) |
| PET foil plasma | 11.0 | 95.9 | 14.107 | 48.0 (0.3) |
| PE foil plasma | 10.9 | 96.1 | 17.522 | 39.0 (0.2) |
| *Planktonic* | | | | |
| CMC (1% w/v) | 64.4 | 99.6 | 3.998 | 64 (1.6) |
| Alginate (1% w/v) | 15.3 | 75.6 | 2.450 | 32 (1.3) |
| Chitin (1% w/v) | 138.4 | 99.59 | 5.276 | 52 (1.0) |
| PET powder (1% w/v) | 153.4 | 99.8 | 5.159 | 69 (1.3) |
| BHET (30 mM) | 20.6 | 94.6 | 3.396 | 140 (4.1) |
| BHET (5.0 mM) | 14.6 | 875 | 7.680 | 1.289 (16.8) |
| BHET (0.5 mM) | 15.9 | 90.2 | 11.850 | 69.0 (0.6) |
| TPA (1mM) | 16.4 | 91.2 | 10.968 | 110.0 (1.0) |
| DMSO | 17.1 | 86.8 | 8.434 | 44.0 (0.5) |
| No additional carbon source added | 127.9 | 99.9 | 6.017 | 44.0 (0.8) |

Cells were grown in artificial sweater medium (ASWM) and in part supplemented with various listed carbon sources. Biofilm cultures were grown in ASWM and planktonic cultures in 1:10 diluted ASWM. PE and PET plasma indicate plasma-treated foil. Data are mean values of three independent biological experiments. Transcriptome data analysis results are available in the supplementary table S1.

**TABLE S3: Significantly differentially expressed genes in PET vs. PE grown biofilms of DSM21264**

| Locus tag | Predicted Function | Log2-foldchange |
|---|---|---|
| AAC977_01250 | predicted glutathione peroxidase, *gene* | 2.73 |
| AAC977_19780 | alkyl hydroperoxide reductase subunit F, *aphF* | 2.41 |
| AAC977_19785 | alkyl hydroperoxide reductase subunit C, *aphC* | 2.33 |
| AAC977_14955 | Dps family protein | 2.02 |

**TABLE S4: Log2-foldchanges of known prophage genes in the genome of DSM 21264 grown in the presence of BHET (0.5, 5.0 and 30 mM) in planktonic cultures.**

| Locus tag | Predicted function | 0.5 mM BHET | 5 mM BHET | 30 mM BHET |
|---|---|---|---|---|
| **VGPH01** | | | | |
| AAC977_19445 | hypothetical protein | 0.25 | 0.29 | -2.12 |
| AAC977_19450 | hypothetical protein | 0.16 | 0.46 | -2.07 |
| AAC977_19455 | putative phage tail protein | 0.33 | 0.44 | 1.02 |
| AAC977_19460 | baseplate J/gp47 family protein | 1.12 | 1.11 | 2.78 |
| AAC977_19465 | phage GP46 family protein | -0.21 | 0.93 | 3.85 |
| AAC977_19470 | phage baseplate assembly protein V | 0.36 | 0.40 | 2.85 |
| AAC977_19475 | phage baseplate assembly protein | 0.18 | 0.50 | 3.03 |
| AAC977_19480 | DNA circularization N-terminal domain-containing protein | 0.33 | 1.60 | 3.65 |
| AAC977_19485 | phage tail tape measure protein | 0.29 | 0.85 | 2.56 |
| AAC977_19490 | phage tail assembly protein | 0.69 | 1.67 | 4.09 |
| AAC977_19495 | phage tail tube protein | 1.40 | 1.99 | 2.49 |
| AAC977_19500 | phage tail sheath subtilisin-like domain-containing protein | 0.72 | 0.09 | 2.33 |
| AAC977_19505 | DUF2635 domain-containing protein | -0.11 | 0.60 | 2.85 |
| AAC977_19510 | hypothetical protein | -0.27 | 1.07 | 2.61 |
| AAC977_19515 | phage virion morphogenesis protein | 1.73 | 0.79 | 4.90 |
| AAC977_19520 | DUF1320 domain-containing protein | -0.17 | -1.28 | 0.31 |
| AAC977_19525 | hypothetical protein | -0.12 | 0.28 | 2.41 |
| AAC977_19530 | hypothetical protein | -0.02 | 0.19 | 3.48 |
| AAC977_19535 | hypothetical protein | 2.93 | 3.36 | 4.36 |
| AAC977_19540 | Mu-like prophage major head subunit gpT family protein | 0.47 | 1.88 | 3.29 |
| AAC977_19545 | phage protease | -0.17 | 1.86 | 5.22 |
| AAC977_19550 | phage minor head protein | -0.04 | 0.06 | 2.75 |
| AAC977_19555 | DUF935 domain-containing protein | 0.25 | 1.18 | 3.72 |
| AAC977_19560 | terminase family protein | 0.95 | 1.91 | 3.93 |
| AAC977_19565 | hypothetical protein | -2.21 | -1.45 | 4.18 |
| AAC977_19570 | DUF3486 family protein | 3.54 | 3.48 | 3.71 |
| AAC977_19575 | ArsR family transcriptional regulator | NA | NA | 5.10 |
| AAC977_19580 | DUF2730 family protein | -0.73 | -0.14 | 6.07 |

| | | | | |
|---|---|---|---|---|
| AAC977_19585 | TraR/DksA C4-type zinc finger protein | 0.00 | 2.48 | 0.00 |
| AAC977_19590 | hypothetical protein | 1.14 | 1.62 | 4.14 |
| AAC977_19595 | hypothetical protein | 0.00 | 3.81 | 7.56 |
| AAC977_19600 | DUF5675 family protein | 2.22 | -0.34 | 6.02 |
| AAC977_19605 | hypothetical protein | -0.17 | -0.37 | 0.80 |
| AAC977_19610 | Mor transcription activator family protein | -2.49 | -1.25 | 3.43 |
| AAC977_19615 | hypothetical protein | -0.72 | -0.64 | -0.44 |
| AAC977_19620 | regulatory protein GemA | 0.07 | 0.26 | -1.08 |
| AAC977_19625 | hypothetical protein | -2.40 | 0.82 | 2.18 |
| AAC977_19630 | hypothetical protein | -0.35 | 1.25 | 1.05 |
| AAC977_19635 | | NA | NA | NA |
| AAC977_19640 | hypothetical protein | -0.09 | -1.14 | 1.88 |
| AAC977_19645 | DUF2786 domain-containing protein | 0.08 | 1.04 | 2.32 |
| AAC977_19650 | hypothetical protein | 1.86 | 1.45 | 3.22 |
| AAC977_19655 | DUF3164 family protein | -1.35 | 1.14 | 2.46 |
| AAC977_19660 | hypothetical protein | 0.00 | 3.01 | 3.73 |
| AAC977_19665 | hypothetical protein | -1.95 | -1.19 | 2.18 |
| AAC977_19670 | hypothetical protein | 0.82 | 2.51 | 4.09 |
| AAC977_19675 | hypothetical protein | -0.14 | 2.83 | 4.26 |
| AAC977_19680 | hypothetical protein | -2.75 | -2.00 | 4.73 |
| AAC977_19685 | AAA family ATPase | -0.26 | 0.45 | 3.30 |
| AAC977_19690 | transposase domain-containing protein | 0.39 | 1.68 | 3.43 |
| AAC977_19695 | helix-turn-helix domain-containing protein | 0.00 | 3.38 | 3.60 |
| AAC977_19700 | S24 family peptidase | -0.04 | -0.42 | -0.46 |
| **VGPH02** | | | | |
| AAC977_20360 | hypothetical protein | 0.43 | 1.99 | 1.15 |
| AAC977_20365 | helix-turn-helix transcriptional regulator | -0.18 | 1.31 | 4.03 |
| AAC977_20370 | hypothetical protein | -0.90 | 0.45 | 3.77 |
| AAC977_20375 | hypothetical protein | -0.28 | 0.95 | 5.06 |
| AAC977_20380 | DNA-binding protein | -0.24 | 0.53 | 5.25 |
| AAC977_20385 | hypothetical protein | 0.40 | 1.35 | 6.31 |
| AAC977_20390 | hypothetical protein | 0.38 | 1.55 | 4.96 |
| AAC977_20395 | phage tail protein | 0.27 | 1.34 | 4.52 |
| AAC977_20400 | hypothetical protein | 0.09 | 0.53 | 5.79 |
| AAC977_20405 | phage tail protein | 0.31 | 1.14 | 5.43 |
| AAC977_20410 | phage tail protein | -0.83 | 1.21 | 5.85 |
| AAC977_20415 | baseplate J/gp47 family protein | 0.20 | 1.21 | 6.81 |
| AAC977_20420 | DUF2590 family protein | -4.26 | -1.30 | 6.56 |
| AAC977_20425 | hypothetical protein | 0.46 | 1.52 | 4.07 |
| AAC977_20430 | hypothetical protein | 0.82 | 3.02 | 8.18 |
| AAC977_20435 | putative phage tail assembly chaperone | -0.88 | 2.98 | 6.89 |
| AAC977_20440 | phage protein | 0.18 | 2.67 | 6.02 |
| AAC977_20445 | DUF2586 domain-containing protein | 0.84 | 3.03 | 5.93 |
| AAC977_20450 | phage tail protein | 1.48 | 2.07 | 3.70 |
| AAC977_20455 | ogr/Delta-like zinc finger family protein | 0.03 | -0.20 | -0.25 |
| AAC977_20460 | hypothetical protein | 3.47 | 3.83 | 5.85 |

| | | | | |
|---|---|---|---|---|
| AAC977_20465 | hypothetical protein | 0.46 | -0.64 | 4.16 |
| AAC977_20470 | lysozyme | -0.61 | -0.43 | 2.82 |
| AAC977_20475 | hypothetical protein | 2.47 | 2.32 | 1.43 |
| AAC977_20480 | hypothetical protein | 0.16 | 0.82 | 4.24 |
| AAC977_20485 | S24 family peptidase | -0.29 | -1.19 | 0.23 |
| AAC977_20490 | anaerobic C4-dicarboxylate transporter | 0.29 | 0.73 | 4.42 |
| **VGPH03** | | | | |
| AAC977_09080 | contractile injection system protein%2C VgrG/Pvc8 family | 0.21 | 0.57 | 0.29 |
| AAC977_09085 | tail protein X | 0.94 | 1.01 | 0.91 |
| AAC977_09090 | phage tail protein | 0.92 | 1.23 | 0.35 |
| AAC977_09095 | phage tail tape measure protein | 0.16 | -0.14 | 0.39 |
| AAC977_09100 | phage tail assembly protein | 0.47 | -0.98 | 0.81 |
| AAC977_09105 | phage major tail tube protein | 0.15 | 1.36 | -2.13 |
| AAC977_09110 | phage tail protein | 0.28 | -0.37 | -1.41 |
| AAC977_09115 | hypothetical protein | -0.04 | -0.56 | -0.57 |
| AAC977_09120 | hypothetical protein | 0.57 | 0.24 | -0.20 |
| AAC977_09125 | phage tail protein | -0.14 | -0.52 | 0.34 |
| AAC977_09130 | hypothetical protein | -0.14 | -0.71 | 2.20 |
| AAC977_09135 | hypothetical protein | 0.35 | -0.10 | 2.04 |
| AAC977_09140 | tail fiber assembly protein | 0.29 | 0.18 | -0.44 |
| AAC977_09145 | hypothetical protein | 0.16 | -0.18 | 0.35 |
| AAC977_09150 | hypothetical protein | -0.04 | -0.21 | 0.40 |
| AAC977_09155 | hypothetical protein | 0.49 | 0.04 | 0.05 |
| AAC977_09160 | hypothetical protein | -0.07 | 0.21 | 1.09 |
| AAC977_09165 | SUMF1/EgtB/PvdO family nonheme iron enzyme | 0.37 | 1.54 | 0.75 |
| AAC977_09170 | phage tail protein | 0.15 | 1.60 | 0.00 |
| AAC977_09175 | phage tail protein I | 0.55 | 0.12 | 1.12 |
| AAC977_09180 | baseplate J/gp47 family protein | 0.45 | 0.09 | -0.34 |
| AAC977_09185 | phage baseplate protein | 0.15 | -0.64 | 0.72 |
| AAC977_09190 | hypothetical protein | 1.47 | 2.24 | 2.86 |
| AAC977_09195 | phage baseplate assembly protein V | 0.80 | 0.51 | 0.92 |
| AAC977_09200 | hypothetical protein | -0.26 | -0.29 | 2.01 |
| AAC977_09205 | hypothetical protein | -0.08 | -0.50 | 3.44 |
| AAC977_09210 | S24 family peptidase | -0.43 | 0.02 | 1.04 |

**Table S5: Confocal microscope settings.**

| **Physical parameter** | **Settings** | |
|---|---|---|
| | Propidium iodide | SYTO 9 |
| Gamma value | 1.0 | |
| Pinhole | 46 µm | |
| Detector digital | 1.0 | |
| Laser intensity | 0.2 % | |
| Laser wavelength | 561 nm | 488 nm |
| Detection wavelength | 560-700 nm | 410-560 nm |
| Detector gain | 663 V | 685 V |
